# Single-molecule multimodal timing of *in vivo* mRNA synthesis

**DOI:** 10.1101/2025.04.27.650906

**Authors:** A.J. Sethi, Marco Guarnacci, Muhammad Bilal, Karthik Subramanian Krishnan, Azusa Hayashi, Madhu Kanchi, Takayuki Nojima, Thomas Preiss, Eduardo Eyras, Rippei Hayashi

## Abstract

mRNA synthesis requires extensive pre-mRNA maturation, the organisation of which remains unclear. Here, we directly sequence pre-mRNA without metabolic labelling or amplification to resolve transcription and multimodal pre-mRNA processing at single-molecule resolution. Using poly(A) tail measurement, we distinguish transcriptionally engaged pre-mRNA from polyadenylated transcripts, revealing that splicing is substantially delayed compared to previous estimates. Splicing is rare for at least 10 kb behind elongating RNA polymerase II such that thousands of genes remain largely unspliced during transcription, revising the notion that splicing is predominantly co-transcriptional. Progressive splicing becomes apparent only on polyadenylated transcripts, suggesting that splicing is commonly activated after 3′ end formation. Unexpectedly, we find abundant m6A on unspliced pre-mRNA, indicating that a substantial portion of RNA methylation precedes splicing. This m6A methylation is suppressed around exon boundaries, indicating that m6A topology is established prior to exon junction complex deposition. Finally, we demonstrate that 3′ end cleavage occurs multiple kilobases behind transcription in coordination with efficient transcription termination, and further highlight recursive 3′ end formation across hundreds of genes. Conserved from human cells to mouse tissues, we illustrate a revised timeline for mRNA synthesis wherein cleavage, termination, and m6A deposition occur earlier than, or at least partially decoupled from, the bulk of splicing, reframing the sequence of early mammalian gene expression.

## Introduction

RNA polymerase II (RNAPII) transcription is coupled with rapid spliceosome assembly across exon boundaries^1–8^. Splicing is thought to occur rapidly after intron transcription, but kinetic estimates vary substantially^1,2,9–20^. RNAPII-bound RNA sequencing suggests that half of introns are rapidly spliced while Pol II is in the next exon, with remaining introns requiring kilobase-scale downstream transcription before splicing^1^. Chromatin-bound and intron lariat RNA sequencing reports that splicing is over 50% complete when RNAPII is less than 160 nt^2^ or 1500 nt^9^ downstream of the intron. In contrast, metabolic labelling indicates a multi-kilobase transcriptional delay before splicing catalysis occurs^10^. Despite converging on co-transcriptional timing, these estimates imply vastly differing organisation and cooperativity between transcriptional and splicing programs.

Splicing regulates mRNA fate via exon-junction complex (EJC) deposition, coordinating adjacent splicing events^21,22^ and inhibiting local N6-methyladenosine (m6A) RNA methylation^23–28^. m6A is proposed to be deposited largely post-transcriptionally^29^, consistent with a model where rapid co-transcriptional splicing directs post-transcriptional m6A topology via EJC placement. Paradoxically, enrichment of metabolically labelled nascent RNA reveals m6A near unspliced exons^30^, and the m6A methylation machinery can interact with the elongating RNAPII^26^, permitting co-transcriptional methylation. How splicing and m6A assemblies cooperate during mRNA synthesis remains elusive due to a paucity of methods to simultaneously assess these processes during transcription^31^.

Here, we map multiple pre-mRNA processing modalities during endogenous transcription, overcoming prior limitations of metabolic labelling, partial fragmentation and amplification. Working in cells and *in vivo*, we demonstrate that splicing catalysis is typically delayed until RNAPII transcribes to a lower bound of at least 10,000 nucleotides beyond intron boundaries, revealing that pre-mRNA is largely unspliced at the time of cleavage and polyadenylation. We demonstrate that m6A accumulates rapidly on unspliced yet polyadenylated pre-mRNA, but is already prohibited at exon boundaries, challenging the role of exon-junction complex as the gatekeeper of m6A topology. Our findings recast current concepts in existing models of mammalian mRNA synthesis, prompting a re-evaluation of the organisation and cooperativity of early eukaryotic gene expression.

## Results

### Development of a multimodal direct pre-mRNA sequencing technique

Identification of co-transcriptional processing events requires discrimination between pre-mRNA undergoing transcription (pre-mRNA^e^) and polyadenylated pre-mRNA undergoing post-transcriptional maturation (pre-mRNA^a^). To address this, we developed dFORCE (direct fractionated observation of RNA coupled to elongation), a single-molecule direct pre-mRNA sequencing technique that simultaneously profiles transcription and pre-mRNA maturation (Fig. 1a). dFORCE leverages the capabilities of direct RNA sequencing to concurrently resolve multiple pre-mRNA processing modalities, including poly(A) tail length^32^, RNA modification^33^, and splicing status, inferring the extent of transcription by directional sequencing from pre-mRNA 3’ ends (Fig. 1b).

**Fig. 1.**
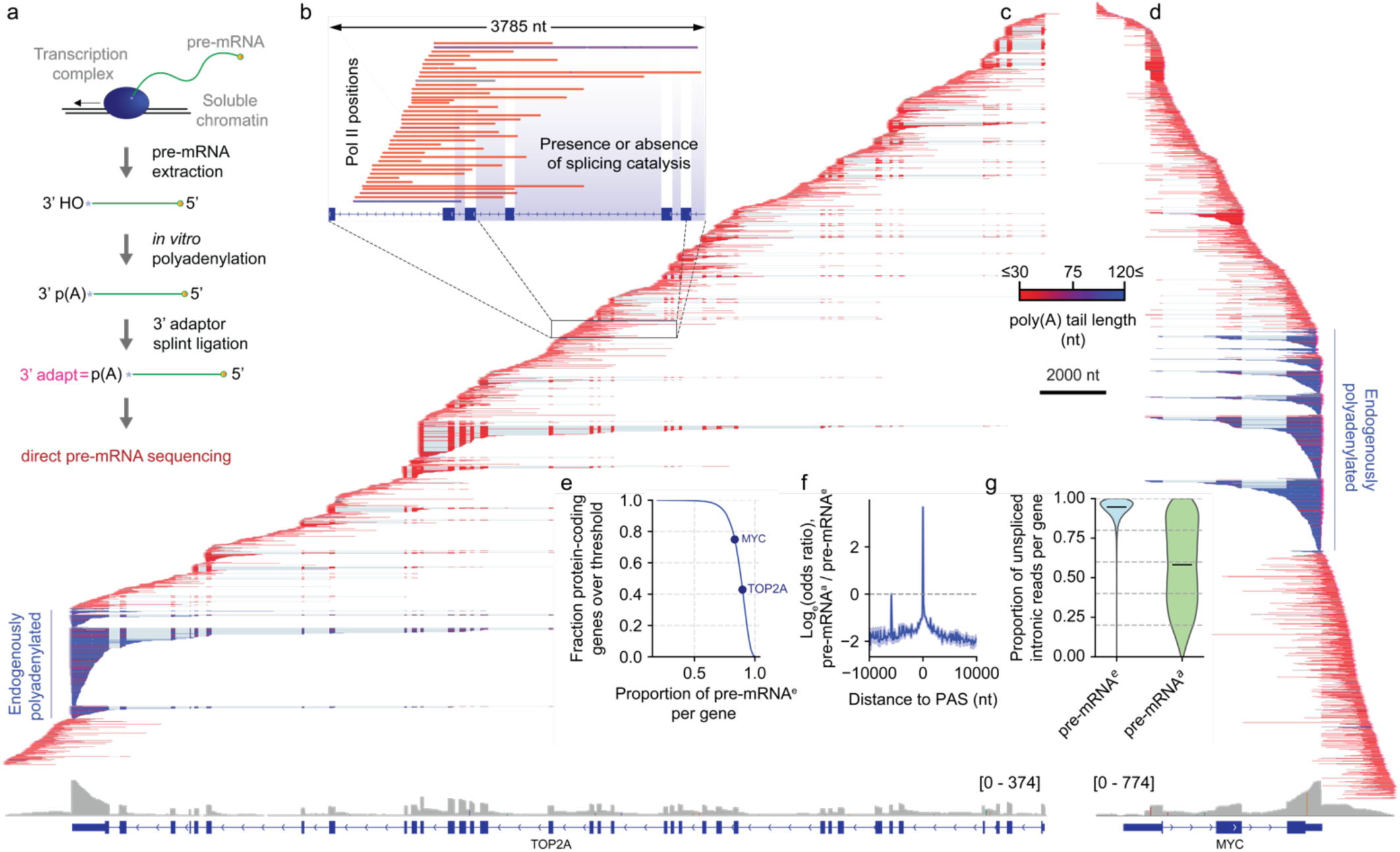
dFORCE separates transcriptionally engaged pre-mRNAe and endogenously polyadenylated pre-mRNAa molecules. **(a)** Strategy of dFORCE, a direct pre-mRNA sequencing technique. Transcriptionally engaged pre-mRNA is isolated from soluble chromatin via polymerase intact nascent transcript (POINT) technology1 and subjected to nanopore direct RNA sequencing. **(b)** Single molecule dFORCE reads capture heterogenous transcription positions with long alignment lengths, enabling the resolution of splicing catalysis against transcription elongation. **(c-d)** IGV browser views of single-molecule dFORCE reads at *TOP2A* and *MYC* loci from representative dFORCE replicates in HeLa cells. Reads are coloured by poly(A) tail length and sorted by alignment 3’ of ends, representing RNA polymerase II position, across the transcription unit. Total coverage over gene regions and annotated gene structures are indicated at the bottom of the figure. Length scale and poly(A) tail length scale are central, shared between the two panels. **(e)** Proportion of co-transcriptional pre-mRNAe reads per protein-coding gene, for 11,614 protein-coding genes with at least 50 poly(A)-classified reads. **(f)** Loge odds-ratio of post-transcriptional pre-mRNAa versus co-transcriptional pre-mRNAe in 100 nt bins of distance between alignment 3’ ends and the nearest annotated poly(A) sites, for 6.8 x 106 protein-coding alignments. Only bins with at least 50 alignments are displayed. **(g)** Proportion of unspliced intron-spanning reads calculated over the total of unspliced intron-spanning and spliced non-intron spanning reads per gene (y-axis), for 10,692 pre-mRNAe and 2,551 pre-mRNAa genes with at least 50 splicing-classified alignments in that poly(A) length classification.

After optimising large-scale RNAPII immunoprecipitation (Fig. S1a), we performed 6 replicates of dFORCE in HeLa cells, generating 24.1 million primary alignments. We also sequenced total mRNA to profile mature mRNA processing. Genome coverage was highly consistent between dFORCE replicates and predominantly spanned protein-coding transcripts despite omission of an rRNA-depletion step, indicating successful RNAPII pulldown (Fig. S1b-c). Protein-coding gene alignments ranged from 80 nt to beyond 12.5 kb in length and covered over 4000 genes with at least 1000 alignments (Fig. S1d-e). Introns and downstream of PAS regions were deeply sequenced, confirming successful isolation of transcriptionally engaged pre-mRNA (Fig. 1c-d).

### dFORCE distinguishes endogenously polyadenylated pre-mRNA

We noticed that poly(A) tails were elongated on alignments ending at poly(A) sites, suggesting that endogenously polyadenylated pre-mRNA^a^ could be distinguished from only *in vitro* polyadenylated pre-mRNA^e^ based on the presence of long poly(A) tails (Fig. 1c-d). To separate these fractions, we normalised pre-mRNA poly(A) tail lengths to distributions observed on endogenously transcribed non-polyadenylated infrastructural transcripts, producing bimodal Z-score distributions (Fig. S1f-g). We classified pre-mRNA with poly(A) length z-scores ≥ 3.6 as endogenously polyadenylated (pre-mRNA^a^) and ≤ 1.8 as non-polyadenylated (pre-mRNA^e^). Across replicates, 90% of pre-mRNA were reproducibly classified as pre-mRNA^e^, whereas around 10% were classified as pre-mRNA^a^ (Fig. 1e and S1h).

If long poly(A) tails in pre-mRNA^a^ reads arose from endogenous cleavage and polyadenylation, we would expect pre-mRNA^a^ alignments to terminate at known mRNA poly(A) sites. Indeed, genome-wide analysis demonstrated that the 3′ ends of pre-mRNA^a^ were enriched at annotated poly(A) sites (odds-ratio = 39.4, p < 10^-300^), indicating that pre-mRNA^a^ tails corresponded to sites of endogenous polyadenylation (Fig. 1f). Further, while 95% of pre-mRNA^e^ reads spanned unspliced introns, only 58% of pre-mRNA^a^ reads did so, indicating a nearly 8-fold increase in splicing catalysis (p < 10^-300^; Fig. 1g). Thus, dFORCE separates non-polyadenylated pre-mRNA^e^ from already polyadenylated but RNAPII affiliated pre-mRNA^a^, enabling distinction of co- and post-transcriptional RNA processing events.

### The landscape of pre-mRNA 3’ polyadenylation

We next applied dFORCE to profile pre-mRNA polyadenylation, detecting 42,000 candidate poly(A) sites (PAS) across 11.6 x 10^3^ coding genes, and recapitulating the canonical <u>AAUAAA</u> poly(A) signal^34^ (Fig. 2a-b). Most of these PAS (79%) were also detected by total mRNA sequencing, 86% were annotated in PolyASite 2.0^35^, and 92% of genes contained at least one PAS coinciding with an Ensembl-annotated site^36^, confirming recovery of genuine poly(A) sites by dFORCE (Fig. S2a-c). Notably, 18% of PAS (7.3 × 10^3^) mapped to introns, indicating widespread intronic polyadenylation with broad metagene distribution (Fig. 2c and S2d). Most intronic PAS occurred downstream of canonical start codons, and over half were also detected in total mRNA, raising the possibility of their translation (Fig. S2e-f). Finally, unannotated splicing events were 3.5-fold enriched at intronic PAS, implying coordination between cryptic splicing and polyadenylation (Fig. S2g).

**Fig. 2.**
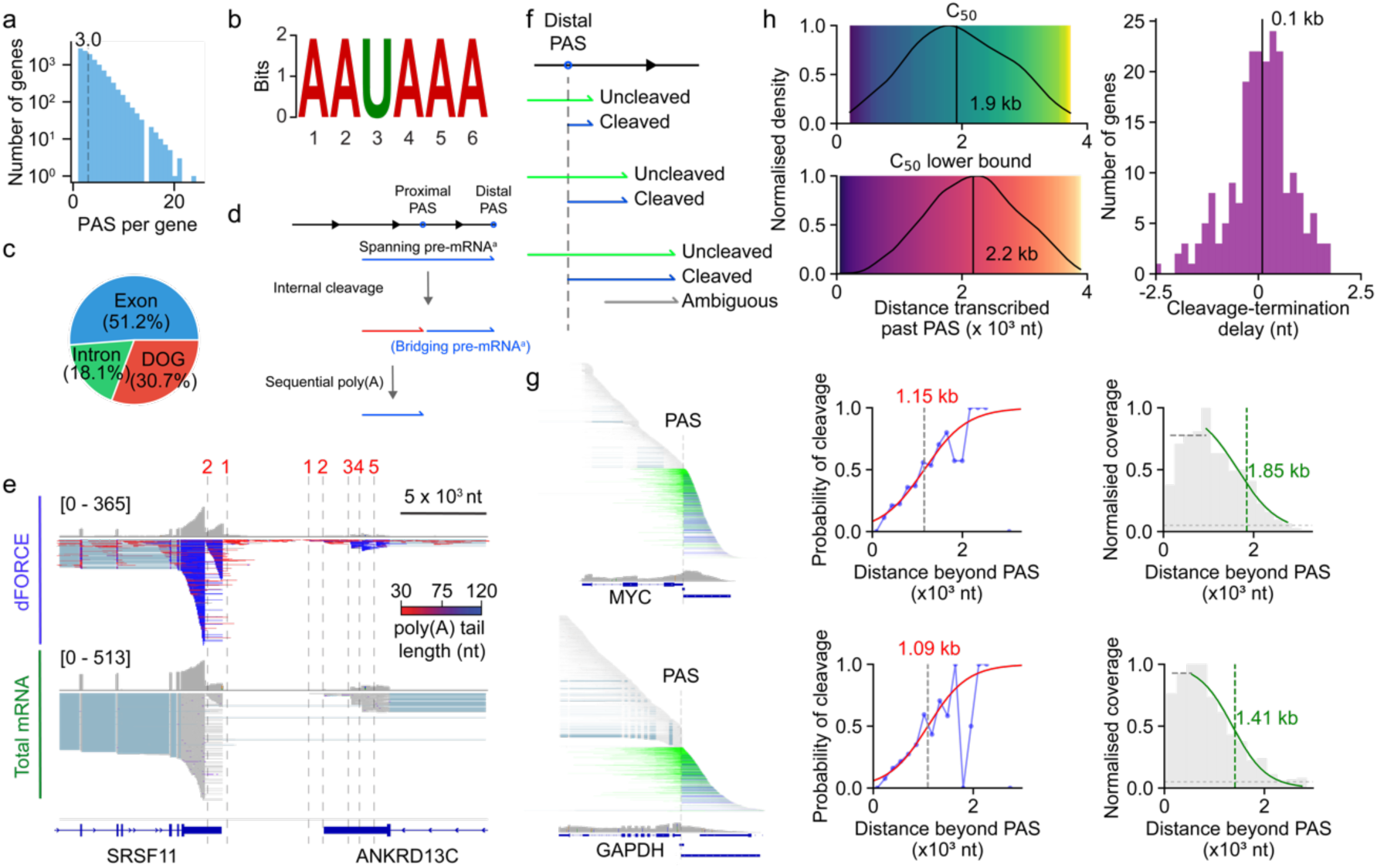
dFORCE resolves the timing of cleavage, polyadenylation and termination at single-molecule resolution. **(a)** Number of consensus poly(A) sites identified by dFORCE (PAS) across 11.6 x 103 protein-coding genes, for 4.2 x 104 total PAS. Dashed line indicates median PAS count per gene. **(b)** The most enriched hexamer sequence motif identified within 40-nucleotide flanking sequences of all PAS sites, discovered using MEME (see methods). **(c)** Distribution of metagene regions (intron, exon, downstream of gene) where PAS were found for sites described in panel (a) **(d)** Schematic indicating the process of recursive polyadenylation. Alignment colours correspond to polyadenylation status with blue alignments being polyadenylated and red alignments not. ’Bridging reads’ are highlighted as the downstream intermediates of recursive polyadenylation. **(e)** Putative recursive polyadenylation sites (indicated with dashed vertical lines) at the 3’ ends of the adjacent *SRSF11* and *ANKRD13C* loci. Poly(A) sites are numbered from most distal to most proximal. Top panel: All dFORCE alignments. Bottom panel: total mRNA alignments. **(f)** Schematic indicating the classification of cleaved, uncleaved and ambiguous reads around distal poly(A) sites. **(g)** Classification of pre-mRNAe alignments and derivation of cleavage C50 and termination T50 values for *MYC* and *GAPDH*. (Left panels) Classification of dFORCE pre-mRNAe alignments that span the cleavage site (green) and alignments that reached near the poly(A) site but did not span it (blue). All other alignments are shown in grey. (Centre panels) Binned cleavage probabilities (blue circles; bin width ∼200 nt) fitted with a logistic regression (red line); dashed grey line indicates the position corresponding to 50% cleavage probability (Right panels) Coverage fraction downstream of PAS shown in 200 nt bins, normalized to peak coverage. Logistic regression (green line) is fitted to the coverage decline; dashed green line marks the position where coverage falls to half the median peak fraction. **(h)** (Upper left): Distribution of C50 distances per gene for 254 genes where cleavage kinetics could be estimated. (Lower left): Distribution of lower bound on C50 distances for 2,867 genes where cleavage was not measurable within our alignment length distribution. Lower bound was derived from the 99% of distance transcribed past the PAS where uncleaved alignments were measured. (right) Density of the difference between median cleavage (C50) and termination (T50) for 221 genes where both values could be estimated. Median values are indicated on all plots.

Intriguingly, we observed recursive polyadenylation, where an already polyadenylated transcript was internally cleaved and subsequently internally polyadenylated^37^. This was indicated by the accumulation of pre-mRNA^a^ with 3′ ends mapping to a distal PAS and 5’ ends mapping to a proximal PAS, i.e., PAS-bridging pre-mRNA^a^ (Fig. 2d). We highlight this pattern at *SRSF3* and *ANKRD13C* loci (Fig. 2e), where bridging patterns suggest internal cleavage and polyadenylation of a pre-mRNA already polyadenylated at the distal PAS. We identified PAS-bridging pre-mRNA^a^ alignments at over 1000 genes. These were a median of 531 nt apart, and affected genes displayed strong enrichment in RNA regulation, including m6A-related roles (Fig. S2h-j). Furthermore, nearly 3% of PAS were bridged to both upstream and downstream PAS, suggesting multiple rounds of recursive polyadenylation (Fig. S2k).

Additionally, we found a localised enrichment of pre-mRNA^a^ 5’ ends at bridged proximal PAS, supporting the notion that the 5’ end of bridging reads arose from internal cleavage of a full-length pre-mRNA^a^ (Fig. S2l). Moreover, for most proximal recursive poly(A) sites, we also detected spanning alignments, i.e. pre-mRNA^a^ ending at the distal PAS that were uncleaved at the proximal PAS, again indicating that polyadenylation at the distal PAS preceded cleavage at the proximal PAS (Fig. S2m). This supports that upstream pre-mRNA^a^ alignments arise from specific cleavage of pre-mRNA^a^ at internal poly(A) sites. Finally, distal bridged PAS were more abundant in pre-mRNA than single PAS (p = 7.9 x 10^-29^, Fig. S2n), consistent with previous reports^37^. Taken together, these results extend previous reports of widespread recursive polyadenylation, confirming that polyadenylation, similar to splicing, can be a multi-step process across hundreds of human genes^29,37^.

### Single-molecule kinetics of cleavage and termination

dFORCE provides a unique opportunity to delineate the timing of co-transcriptional 3’ cleavage and transcription termination by analysing the properties of non-polyadenylated reads (pre-mRNA^e^) extending beyond distal poly(A) sites. We classified these alignments as cleaved or uncleaved, based on their 5’ ends, and measured how the frequency of cleaved alignments changed with increasing transcription beyond the PAS (Fig. 2f). From this data, we modelled C_50_, the distance beyond the PAS at where cleavage reached 50% efficiency, and T_50_, the distance where the density of RNAPII active sites dropped to half the coverage downstream of the PAS, implying transcription termination. We illustrate these measurements at *MYC* and *GAPDH* loci, where coverage was sufficient to estimate cleavage and termination transcriptional kinetics (Fig. 2g).

For the 254 protein-coding genes where 3’ cleavage reached 50% efficiency within observed alignment length distribution, the mean C_50_ distance lay 1.9 x 10^3^ nt downstream of the PAS (Fig. 2h). For an additional 2,687 genes, the C_50_ distance was found outside our read length distribution. However, using observed alignment lengths, we estimated a lower bound on C_50_ to be 2.1 x 10^3^ nt beyond the PAS (Fig. 2h; bottom). This suggests that 3’ cleavage, which precedes polyadenylation, is typically delayed by 2000 nt or more following transcription of the poly(A) site.

To infer the extent of transcription before RNAPII termination, we modelled the distance transcribed beyond the PAS using logistic regression of Pol II active site density. Across 2.6 x 10^3^ protein-coding genes, the median T_50_ distance was 2.3 x 10^3^ nt beyond the distal poly(A) site (Fig. S2o). For the 221 genes where we could estimate both a cleavage and termination distance, we compared the difference in these values, finding that transcription termination was delayed relative to cleavage by a median distance of only 95 nt (Fig. 2j). Our analyses with dFORCE provides a first long-read RNA-centric timing of cleavage and termination, indicating that transcription termination occurs in close temporal proximity to cleavage.

### Discrepant evidence for co-transcriptional splicing catalysis

Next, we examined the extent of splicing catalysis during transcription, i.e. co-transcriptional splicing. We found global discrepancies between pre-mRNA^e^ alignments ending in exons, which displayed frequent splicing of upstream introns, and pre-mRNA^e^ alignments ending in downstream introns, where the same splicing events were typically abolished. For instance, *MYC* intron 1 was often spliced for alignments ending in exons 2 and 3 but was consistently unspliced for alignments ending in intron 2 or beyond the PAS (Fig. 3a). Similar patterns were observed at more complex genes, e.g. *TOP2A*, where 35 exons harboured a median of 50% spliced pre-mRNA^e^, versus 0% for adjacent introns (Fig. S3).

**Fig. 3.**
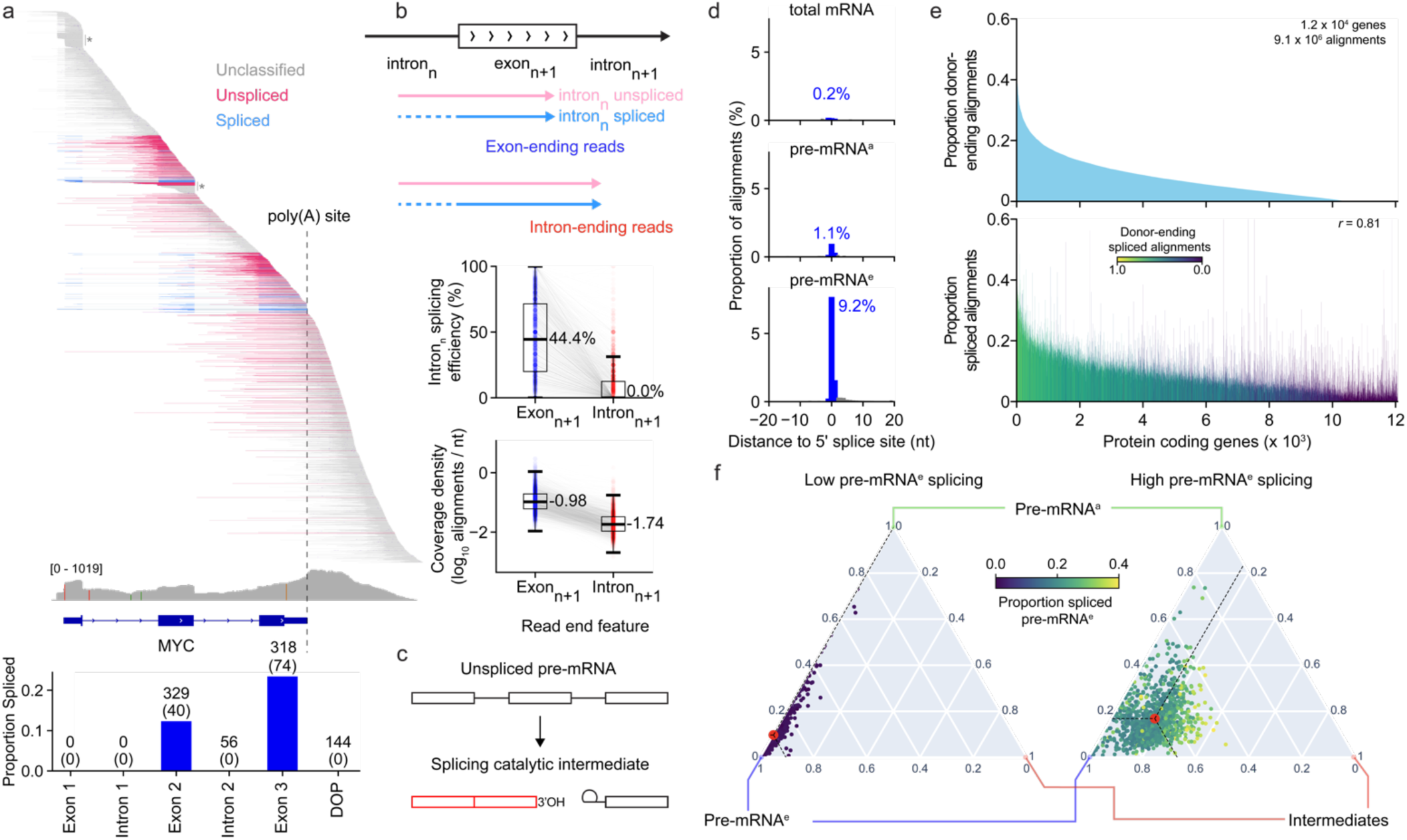
Discrepant evidence for co-transcriptional splicing catalysis. **(a)** Top: Co-transcriptional pre-mRNAe alignments at *MYC*. Alignments are coloured by splicing classification and sorted by mapped 3’ end coordinate. ‘Spliced’ reads contain ligated exon-exon junctions, unspliced reads contain at least one fully transcribed intron and no splice junctions, and all other reads, e.g. those fully contained in a single exon or intron, are unclassified. Only pre-mRNAe alignments longer than 200 nt that lacked cryptic splice junctions are shown. Bottom: Proportion of pre-mRNA*e* reads that are spliced (y-axis) ending at each of the *MYC* gene region (x-axis). Numbers above bars indicate the number of tested and spliced (in parenthesis) reads ending within each exon, intron, or downstream of the poly(A) site (DOP) region. **(b)** Top: Diagram describing the intron-exon-intron splicing test. Reads ending in the downstream exonn+1 and intronn+1 are tested for the proportion of splicing seen in the upstream intronn. Only alignments that span the 5’ boundary of exonn+1 are tested. Middle panel: Proportion of intronn splicing seen in reads ending in the downstream exon and intron, across 40,565 intron-exon-intron trios in the human genome. Lower panel: Total density of 3’ end alignment terminating in same set of exons and introns exonn+1 and intronn+1. The density was calculated as the number of 3’ ends over the feature width. **(c)** Diagram illustrating the production of spliced, non-polyadenylated RNA molecules from 5’ fragment of first step splicing catalytic intermediates. **(d)** Proportion of alignments for total mRNA (upper), dFORCE pre-mRNAa (middle) and dFORCE pre-mRNAe (lower) ending at different positions relative to 126,954 unique 5’ splice site detected in dFORCE. Positions within 1 nt of 5’ splice sites are shaded in blue. Percentage of alignments ending within 1 nt of a 5’ splice site are labelled on the plot in blue. **(e)** Top: Ranking (x-axis) of protein-coding genes by the proportion of pre-mRNAe reads end within 1 nt of a 5’ splice site, classified as catalytic intermediates (y-axis). Bottom: In the same ranking, proportion of pre-mRNAe alignments that display a splice junction (y-axis) across the same set of genes. Genes are coloured by the proportion of spliced reads that end within 1 nt of a 5’ splice site, compared to the number of spliced reads (‘Donor ending spliced alignments). **(f)** Ternary plot representing the proportion of pre-mRNAa, pre-mRNAe (excluding donor-ending reads) and intermediates (i.e. donor-ending pre-mRNAe reads) for protein-coding genes with at least 50 alignments and up to 40% spliced pre-mRNAe. Left panel: bottom 10% of genes according to spliced pre-mRNAe level. Right panel: top 10% of genes. Red dots indicate the average proportions of (non-intermediate) pre-mRNAe, catalytic intermediates, and post-transcriptional pre-mRNAa in each gene set. 1.2 x103 genes are displayed per panel. Middle colour scale indicates the proportion of pre-mRNAe reads that are spliced in each panel. Genes that display more co-transcriptional splicing have more splicing catalytic intermediates and pre-mRNAa, indicating that these are the likely source of observed splicing events. In contrast, genes with little co-transcriptional splicing mainly present only pre-mRNAe as opposed to pre-mRNAe or CI*.

To globally quantify this splicing discrepancy, we systematically compared the splicing efficiency of individual upstream introns (intron_n_) between pre-mRNA^e^ alignments ending either in the next exon (exon_n+1_) or the next intron (intron_n+1_). Testing 40,565 intron-exon-intron trios across 7,698 protein-coding genes, we found that alignments ending in exon_n+1_ displayed frequent splicing of intron_n_ (median 44.4% spliced), but this splicing was acutely lost on alignments ending in intron_n+1_ (median 0.0% spliced; Fig. 3b). Thus, this discrepancy was globally prevalent across the trancriptome. Further, alignment density was a median of 5.7-fold increased across the set of exon_n+1_ versus intron_n+1_ (Fig. 3b). Together, these splicing discrepancies and alignment density patterns were preserved across intron sets of varying length, GC content, and nuclear speckle association^38^ (Fig. S4a-c).

To exclude potential methodological or sample-specific artefacts, we repeated dFORCE in an independent human cell line (K562), and further reanalysed data from previous nascent RNA sequencing experiments^1,10^ (Fig. S4d-e). All exhibited the same phenomenon, with prevalent spliced pre-mRNA ending in exons that was lost on alignments ending in the downstream intron. Under true co-transcriptional splicing catalysis, we would expect that the splicing efficiency of upstream introns should remain constant or increase as RNAPII elongates through downstream regions. Hence, the acute transcriptome-wide loss of spliced pre-mRNA^e^ ending in introns suggests that the majority of spliced pre-mRNA^e^ ending in upstream exons were not transcriptionally engaged and did not represent co-transcriptional splicing.

### Catalytic intermediates inflate co-transcriptional splicing estimates

We found that overrepresentation of pre-mRNA^e^ density within exon_n+1_ correlated strongly with the presence of pre-mRNA displaying splicing of intron_n_ (*R*^2^ = 0.94 across groups; Fig. S4f). We therefore hypothesised that 3’ ends of spliced pre-mRNA^e^ found in exons may not represent the site of transcription, but rather other events. For instance, the 1^st^ splicing step produces upstream catalytic intermediates that end at 5’ splice sites and could harbour previously spliced upstream introns (Fig. 3c). Lacking poly(A) tails, these species would be classified as pre-mRNA^e^ by our method.

First, we estimated the abundance of 5’ catalytic intermediates (CI*) by measuring pre-mRNA^e^ 3’ end density around 5’ splice sites^2,9,10,39^. Nearly 10% of all pre-mRNA^e^, including over half of all spliced pre-mRNA^e^, terminated within 1 nt of a 5’ splice site, indicating that CI* were abundant and a major source of spliced pre-mRNA^e^ (Fig. 3d; Fig. S4g-h). To understand how CI* alignments influenced splicing estimates, we ranked genes by proportions of CI* pre-mRNA^e^, revealing a striking correlation against proportions of spliced pre-mRNA^e^ (*r* = 0.81; Fig. 3e). Finally, excluding CI* pre-mRNA^e^ resulted in substantial correction of exon_n+1_ vs intron_n+1_ splicing discrepancies to a difference of 25% (Fig. S4i). We demonstrate that most spliced pre-mRNA^e^ correspond to splicing catalytic intermediates rather than elongating pre-mRNA, substantially confounding co-transcriptional splicing estimates (Fig. 3f).

### Elongating pre-mRNA is globally unspliced for thousands of nucleotides

To quantify the true extent of co-transcriptional splicing, we assessed two transcriptional windows where 5’ catalytic intermediates or other mRNA fragments would minimally affect our observations. First, during transcription of the last intron (excluding CI* formed at upstream 5’ splice sites), and second, after the transcription of poly(A) sites but before cleavage and polyadenylation (excluding CI* or any fragmentation products of pre-mRNA^a^; Fig. 4a). We classified the splicing of pre-mRNA^e^ ending in each window based on their observed upstream introns, contextualising this with a gene-specific context length corresponding to the maximum aligned distance observed per gene in each window.

**Fig. 4.**
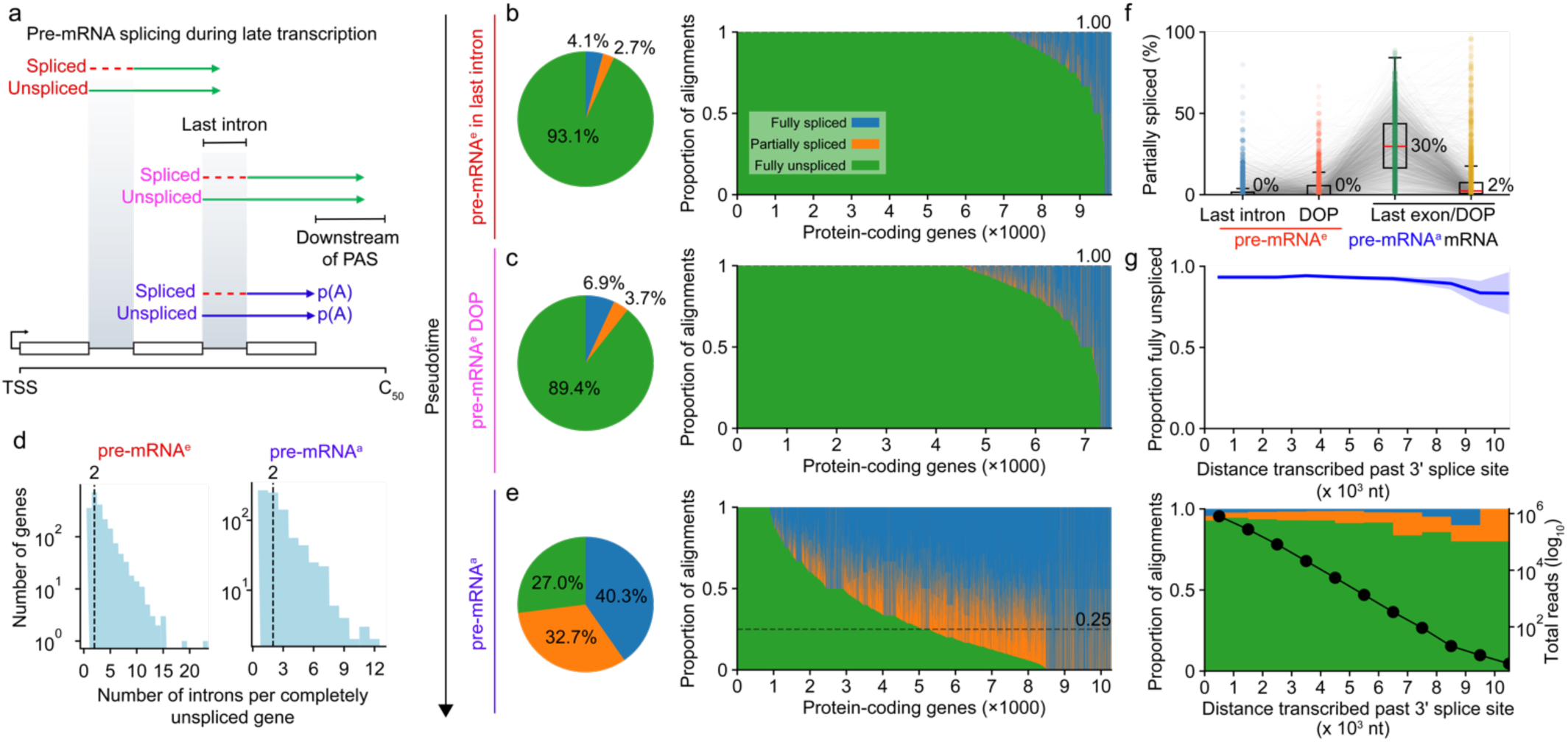
Pre-mRNA splicing catalysis is rare during transcription. **(a)** Schematic outlining analysis of splicing catalysis during transcription through last introns, downstream-of-poly(A) sites (DOP), and following cleavage and polyadenylation. **(b)** Distribution of spliced, unspliced and partially spliced alignments over 9,825 protein-coding genes with pre-mRNAe ending in the last intron that could be assessed. (Left): Pie chart showing distribution of splicing state across all classified pre-mRNAe in the last introns. (Right): Distribution of the same splicing states stratified by gene, where genes are ranked by their proportion of fully unspliced alignments. The dashed horizontal line represents the median proportion of fully unspliced alignments per gene. Green: fully-unspliced read, orange: partially spliced alignment, blue: fully spliced alignment. This classification only considers upstream introns and ignores the splicing of the last intron, which is partially transcribed. **(c)** Similar to (b), for pre- mRNAe ending downstream of the dominant poly(A) sites, for 7,526 protein-coding genes. **(d)** Distribution of intron count for genes which presented completely unspliced alignments (i.e. alignments that simultaneously spanned every intron in the gene). Left: 2,051 protein-coding genes that present completely unspliced alignments in pre-mRNAe. Right: 824 genes that present completely unspliced alignments in pre- mRNAa. Vertical dashed lines indicate median intron counts across gene sets. **(e)** Similar to (b), (c), for protein-coding genes presenting pre- mRNAa ending in the last exon or downstream of the distal poly(A) site. 10,286 genes are plotted, of which 895 only present unspliced alignments. **(f)** Proportion of partially-spliced alignments compared to partially-spliced, fully-spliced and fully unspliced, for 2,542 protein-coding genes with at least 5 assessed alignments in all of pre-mRNAe in the last intron, pre-mRNAe downstream of the poly(A) site, pre-mRNAa in the last exon or downstream of the poly(A) site, and total mRNA ending in the last exon or downstream of the poly(A) site. **(g)** (Top) Smoothed proportion of fully unspliced reads versus extent of transcription past the 3’ splice site, binned in 1000 nt intervals. Shaded band indicates the 95% CI derived from the binomial variance within each rolling window. (Bottom) Stacked bar plot (left y-axis) displaying the proportion of fully unspliced, partially spliced and fully spliced alignments within each bin indicated in the top panel. Colours correspond to panel (b). The overlaid black line (right y- axis) indicates the number of alignments tested within each bin. Alignments assessed from 1.2 x 104 genes are included in this plot. The x-axis (shared) corresponds to the distance between the 3’ splice site and the alignment 3’ end. Note, alignments ending in exons are excluded from this plot (see. Fig. S5f).

First, we assessed upstream splicing during transcription of last introns. Testing 9,825 genes, we recapitulated our previous findings: pre-mRNA^e^ ending within last introns were overwhelmingly (93.1%) fully unspliced across all observed upstream introns (Fig. 4b), to a median context of 2.0 x 10^3^ nt (95% range: 0.4 - 5.2 × 10^3^ nt), resolving 2 introns per gene (95% range: 2 – 7 spanned introns; Fig. S5a). For 72% of genes, all observed pre-mRNA^e^ was fully unspliced to a median context of 1.8 x 10^3^ nt (95%: 0.4 – 4.8 × 10^3^ nt; Fig. S5a). To ensure these results were not unique to last introns, we repeated this analysis over all internal introns, finding similarly, that 92.3% of alignments were fully unspliced to a median context of 3.5 x 10^3^ nt per gene across 10,599 genes (Fig. S5b).

We next measured splicing on uncleaved pre-mRNA^e^ ending downstream of distal poly(A) sites. Assessing 7,526 genes, we found that the majority (89.4%) of pre-mRNA^e^ was fully unspliced (Fig. 4c), to a median context of 3.4 x 10^3^ nt (95%: 1.1 – 6.4 x 10^3^ nt), resolving 2 introns per gene (95%: 1 – 8 spanned introns; Fig. S5c). Over 60% of genes (4,536 genes) presented exclusively unspliced alignments in this region, with a median context length of 3.1 x 10^3^ nt (95%: 1.0 – 6.2 x 10^3^ nt). Overall, we identified 2,051 genes where every intron in the gene was spanned by at least 1 unspliced pre-mRNA^e^, indicating that the primary transcript was entirely unspliced during late transcription (Fig. 4d). These genes harboured up to 23 introns (Fig. S5e) and were spanned by a median of 6 completely unspliced alignments, reinforcing that rapid co-transcriptional splicing catalysis was exceedingly uncommon in the vicinity of elongating RNAPII.

### Splicing catalysis is deferred until after polyadenylation

Given that splicing was deferred during transcription, we next asked whether it was efficient after polyadenylation. To address this, we assessed pre-mRNA^a^, encompassing both newly polyadenylated transcripts, and older transcripts which had persisted on the chromatin. Surveying the last exon and downstream of poly(A) site (DOP) regions of 10,286 protein-coding genes, we found that pre-mRNA^a^ ending in these regions were globally 27% fully unspliced, 33% partially spliced, and 40% fully spliced (Fig. 4e), to a median context of 3.2 x 10^3^ nt (95%: 0.7 – 6.4 x 10^3^ nt), typically resolving 1 unspliced intron per gene (95%: 0 – 7 introns; Fig. S5d). This distribution confirmed that splicing was abundant on chromatin, but only following polyadenylation, rather than during elongation.

Notably, for 895 genes, we observed exclusively unspliced pre-mRNA^a^, indicating that polyadenylation always preceded splicing within the genomic context we could observe. Further, for 824 multi-exon genes, we identified fully unspliced pre-mRNA^a^ alignments that spanned every intron in the gene, indicating that polyadenylation preceded all splicing anywhere in the gene body (Fig. 4d). These measurements are likely underestimates, since sequencing depth was around 10-fold lower for pre-mRNA^a^ vs pre-mRNA^e^. Taken together, these data strongly suggest that splicing catalysis generally commences only after 3′ end formation.

Next, we quantified the abundance of partially spliced RNA as a measure that splicing catalysis was underway. Comparing a set of 2,542 genes with sufficient coverage across fractions, a median of 0% of alignments per gene was partially spliced when RNAPII was in the last intron or downstream of the poly(A) site. Similarly, partial splicing in total mRNA was minimal (median 2%), likely reflecting retained introns. In contrast, partial splicing was enriched on polyadenylated pre-mRNA^a^ (median: 30% per gene), indicating that the intermediates of sequential splicing events could be faithfully detected in our data, but only following cleavage and polyadenylation (Fig. 4f).

### A lower bound for co-transcriptional splicing catalysis

Since dFORCE alignments were typically shorter than 10,000 nt, we could not directly resolve upstream splicing rate during deep transcription within long introns. To estimate this, we pooled non-exon ending pre- mRNA^e^ reads across protein-coding genes and calculated the splicing status of the upstream intron as a function of distance transcribed beyond 3’ splice sites. Fully unspliced pre-mRNA remained dominant throughout this range, up to the maximum measured distances beyond 10^4^ nt (Figs. 4g and S5f). Thus, co-transcriptional splicing is substantially delayed compared to prior estimates, commonly by over ten thousand nucleotides behind transcription.

### The quantitative landscape of pre-mRNA m6A

We leveraged the ability of direct RNA sequencing to resolve RNA modifications at single molecule resolution^33^ to interrogate the timing of m6A deposition during mRNA synthesis. First, we basecalled m6A on total mRNA (Fig. 5a), mapping 1.82 x 10^5^ m6A sites mainly (81.1%) in the known DR<u>A</u>CH sequence motif^40^ (Fig. 5b). Comparison with GLORI^41^ showed strong m6A concordance (*R^2^ =* 0.80), and we recovered known topological features, including exon-exon junction proximal m6A depletion^33^, further validating our sites (Fig. S6a-d). We then basecalled m6A on pre-mRNA, identifying 2.19 x 10^5^ m6A sites. While m6A was abundant on exonic regions of pre-mRNA, it was 17-fold depleted across introns (Figs. 5c and S6e). Further, while exonic m6A was most depleted at exon-intron boundaries, intronic m6A was greatest near boundaries, showing suppression in deeper regions (Figs. 5d and S6f).

**Fig. 5.**
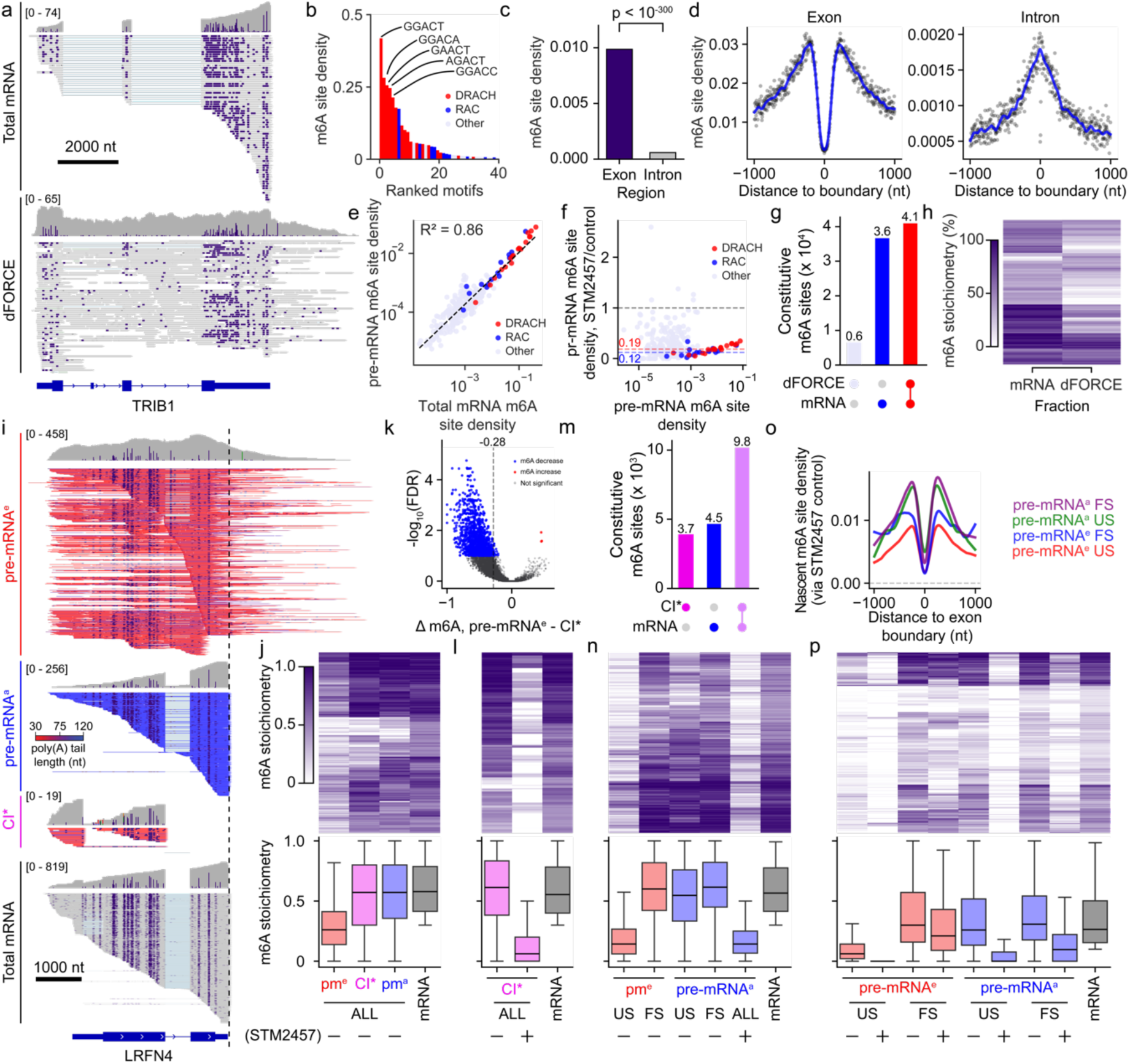
The quantitative landscape of pre-mRNA m6A methylation. **(a)** m6A modifications in total mRNA (upper panel) and pre-mRNA (lower panel) at the *TRIB1* locus. Individual m6A calls are indicated as purple bases. The proportion of methylated reads at each genomic position is indicated in the coverage tracks for each panel. **(b)** Identified m6A sites grouped by A-centred 5-mer (NN<u>A</u>NN) from total mRNA. The y axis (m6A site density) indicates the proportion of genomic instances in which that 5-mer is classified as m6A (≥5% m6A stoichiometry and ≥2 m6A- modified reads) over the total number of genomic instances with that 5-mer that were tested in total mRNA. Only the top 40 5mers are shown. Bars are coloured by motif type, where red corresponds to DR<u>A</u>CH motifs, blue to non-DRACH R<u>A</u>C motifs, and grey to other motif types. **(c)** pre-mRNA m6A site density across exons and intronic features. Density was calculated based on grouping sites by exonic or intronic region. Chi-square test was used. **(d)** Pre-mRNA m6A site density on exons (left panel) and introns (right panel) based on distance to splice sites used in total mRNA. For exons, sites along the x-axis were selected on either side of adjacent exon-exon junctions. For introns, sites at >0 positions are those downstream of a 5’ splice site, whereas <0 positions are those upstream of a 3’ splice site. **(e)** Motif-level NN<u>A</u>NN m6A site density compared between total mRNA (x-axis) and dFORCE pre-mRNA (y-axis) fractions. Density calculations and colour scale correspond to panel (b). **(f)** Scatter plot of m6A site density versus treatment effect for A-centred 5-mers. The x-axis shows the m6A site density of each A-centred 5-mer in dFORCE pre-mRNA, and the y-axis gives the fold-change in m6A density upon STM2457 treatment relative to control. Colour coding and density calculations correspond to panel (c). **(g)** UpSet plot representing the distribution of m6A sites in dFORCE only (light blue), total mRNA only (dark blue), or both (red), for 83,295 constitutive m6A sites (methylated at ≥ 30% stoichiometry in either dFORCE or total mRNA) which were tested in both fractions (≥ 2 reads in both fractions). **(h)** Heatmap of m6A stoichiometry values in total mRNA (left) and dFORCE (right) for 64,356 common sites methylated at ≥ 30% in either fraction with at least 5 reads in each fraction. Sites shown are a subset of those in (h) with higher coverage. **(i)** dFORCE (top 3 panels) and total mRNA (bottom panel) m6A-basecalled alignments at the *LRFN4* locus. dFORCE alignments are coloured by poly(A) tail length and separated into pre-mRNAe, pre-mRNAa and splicing catalytic intermediate (CI*) fractions. Only positive strand alignments are displayed. m6A sites are displayed as in panel (a). **(j)** Top: Heatmap of m6A stoichiometry values for 2,175 m6A sites that are constitutive in total mRNA (≥ 30% stoichiometry) and are tested (i.e. are measurable) in at least 5 alignments in total RNA and each dFORCE fractions. Bottom: Boxplots representing the stoichiometry distribution in the same fractions. pme: pre-mRNAe, CI*: splicing catalytic intermediates, pma: pre-mRNAa. **(k)** Differential m6A methylation test for 9,481 m6A sites detected to be at least 30% methylated in dFORCE pre-mRNAe or CI* and spanned by at least 5 reads in both fractions. x-axis: Difference in site stoichiometry between pre-mRNAe – catalytic intermediates, y-axis: FDR-adjusted p-value. Vertical dashed line indicates the median stoichiometry difference across the sites; Sites hypomethylated on pre-mRNAe (FDR ≤ 0.1 and Δm6A ≥ = -0.1) are coloured in blue, and those hypermethylated on pre-mRNAe (FDR ≤ 0.1 and Δm6A ≥ = 0.1) are coloured in red. **(l)** Similar to (j), comparing n = 1,968 m6A constitutive sites in total mRNA (≥ 30% stoichiometry) that were observed in splicing intermediates (CI*) before and after treatment with STM2457. Layout and colours correspond to (k). **(m)** UpSet plot showing identification of 18,091 constitutive m6A sites detected in pre-mRNA splicing intermediates (CI*) only, total mRNA only, or both fractions. **(n)** Similar to (j,l), testing n = 1,204 sites that were detected as constitutive in total mRNA and tested in at least 5 alignments in all fractions shown. Pre-mRNA fractions are stratified by splicing status (US: unspliced, FS: fully spliced) and by STM2457 treatment status. **(o)** Similar to (d). Plot shows m6A methylation rates on exons on fully spliced and fully unspliced pre-mRNAe and pre-mRNAa fractions, normalised to rates seen following STM2457 treatment for 2 hours. Normalised values indicate the true rates of m6A methylation on these fractions in the prior 2-hour timespan. **(p)** Similar to (k,l,n), testing n = 432 m6A sites identified as being at least 10% m6A methylated in total mRNA, and sequenced by at least 4 alignments in all fractions shown. Groups are stratified by splicing and STM2457 treatment as described in (n).

Preferential pre-mRNA DRACH methylation suggested METTL3 catalysis (Fig. 5e). We validated this by briefly applying STM2457, a potent METTL3 inhibitor, achieving pre-mRNA m6A depletion at DR<u>A</u>CH (81% reduction) and R<u>A</u>C (88% reduction) sites (Fig. 5f). To determine the fate of pre-mRNA m6A, we compared methylation between untreated pre-mRNA and total mRNA. Testing 8.3 x 10^4^ constitutive m6A sites (sites with ≥30% stoichiometry), we found constitutive methylation to be mostly common in both fractions or unique to total mRNA, but rarely specific to pre-mRNA (Fig. 5g-h). Further, strong DR<u>A</u>CH sites silenced in total mRNA were typically already silenced in pre-mRNA (Fig. S6g). Thus, once installed on pre-mRNA, m6A is efficiently displayed in total mRNA.

### Single-molecule timing of m6A deposition

Splicing is thought to determine m6A topology via EJC-mediated steric exclusion, a model which requires m6A deposition to occur after splicing catalysis^29^. Leveraging our isoform-resolved quantitative m6A maps, we revisited this model. We separated m6A across co-transcriptional (pre-mRNA^e^), post-transcriptional (pre- mRNA^a^) and catalytic intermediate (CI*) fractions, producing m6A-resolved mRNA biogenesis maps, illustrated at *LRFN4* (Fig. 5i). Using these maps, we assessed methylation dynamics of constitutive total mRNA sites across stages of mRNA synthesis. While median m6A levels were more than 50% depleted in pre-mRNA^e^ versus total mRNA, they were almost identical between the remaining fractions (Fig. 5j). This suggests that m6A deposition starts co-transcriptionally and pre-mRNA reaches close to cytoplasmic m6A methylation levels prior to the completion of splicing.

To confirm that m6A was enriched on CI*, we performed differential methylation analysis, finding 1,712 sites significantly hypermethylated on CI* versus pre-mRNA^e^ (FDR ≤ 0.1), but only 3 hypomethylated sites (Fig. 5k). We mapped 29,188 m6A sites on CI* corresponding to 5,010 genes (Fig. S6h). While m6A on catalytic intermediates was nearly identical compared to total mRNA at constitutive sites, CI* m6A was ∼90% abolished after STM2457 treatment (Fig. 5l). Of 18,000 constitutive total mRNA sites tested on CI*, over half were constitutively methylated on CI*, corresponding to m6A at 2,450 genes (Fig. 5m). Therefore, we establish that m6A is already deposited before splicing completes at many thousands of genes. Moreover, in addition to the presence of upstream splicing events and 3’ termination at 5’ splice sites, we show that CI* can be distinguished from pre-mRNA^e^ based on m6A hypermethylation alone.

### m6A deposition precedes global splicing catalysis

Given the abundant m6A found on catalytic intermediates, we next asked whether m6A was installed before splicing begins. To address this, we assessed m6A levels across spliced and unspliced pre-mRNA fractions. Despite m6A depletion on fully unspliced (US) pre-mRNA^e^, m6A levels were consistent between spliced pre- mRNA^e^, unspliced or spliced pre-mRNA^a^, and total mRNA (Fig. 5n). These patterns suggested that splicing catalysis was entirely preceded by substantial m6A methylation to near cytoplasmic levels. To validate m6A detected on unspliced pre-mRNA^a^, we compared to STM2457-treated pre-mRNA^a^, finding that m6A was largely abolished to unspliced pre-mRNA^e^ levels. Given that pre-mRNA^a^ is endogenously polyadenylated, our results indicate that constitutive m6A sites are almost entirely methylated after polyadenylation but before splicing begins.

Since we observed m6A before splicing catalysis, and therefore prior to EJC deposition, we next tested whether exon boundary m6A silencing was detected in unspliced pre-mRNA. We examined m6A density around exon boundaries in both spliced and unspliced pre-mRNA, normalised to STM2456 control to identify m6A sites turned over within a 2-hour window (Fig. 5o). Globally, m6A levels were minimal in unspliced pre- mRNA^e^, moderately elevated in spliced pre-mRNA^e^, but markedly high and nearly identical in both fully spliced (FS) and unspliced (US) pre-mRNA^a^. Notably, exon-boundary m6A depletion was consistently observed across all fractions, indicating that these boundary-proximal sites are silenced well before splicing catalysis, and therefore, prior to EJC deposition, likely via alternative mechanisms.

We expanded the analysis including weaker, non-constitutive sites, by analysing RNA sites methylated with ≥10% stoichiometry across multiple splicing and transcriptional fractions. We again observed low co-transcriptional stoichiometries on unspliced pre-mRNA^e^, which were abolished by STM2457 (Fig. 5p), confirming that even modestly methylated sites are installed co-transcriptionally. In contrast, both US and FS pre-mRNA^a^ showed m6A levels similar to total mRNA, underscoring a mature state of methylation that was largely independent of splicing completion (Fig. 5p). Meanwhile, fully spliced pre-mRNA^e^ often carried near-mature methylation, supporting an association between m6A methylation and pre-mRNA maturation.

### dFORCE resolves the *in vivo* sequence of mRNA synthesis

Having established dFORCE in cultured cells, we next interrogated mRNA synthesis under native physiological constraints in the context of mammalian tissue. After optimising chromatin isolation, we applied dFORCE to entire mouse brains, producing 40 million alignments across three consistent replicates (Fig. 6a and S7a). Brain dFORCE profiles closely resembled those observed in HeLa cells (Fig. 6b), displaying strong correlation (*R* = 0.69) of pre-mRNA^e^/pre-mRNA^a^ proportions across pairs of orthologous genes (Fig. S7b-c). dFORCE successfully recovered annotated mouse poly(A) sites for over 90% of 1.3 x 10^4^ tested mouse coding genes, confirming accurate separation of polyadenylated pre-mRNA^a^ (Fig. S7d). Thus, dFORCE successfully isolates pre-mRNA^e^ and pre-mRNA^a^ fractions under *in vivo* conditions.

**Fig. 6.**
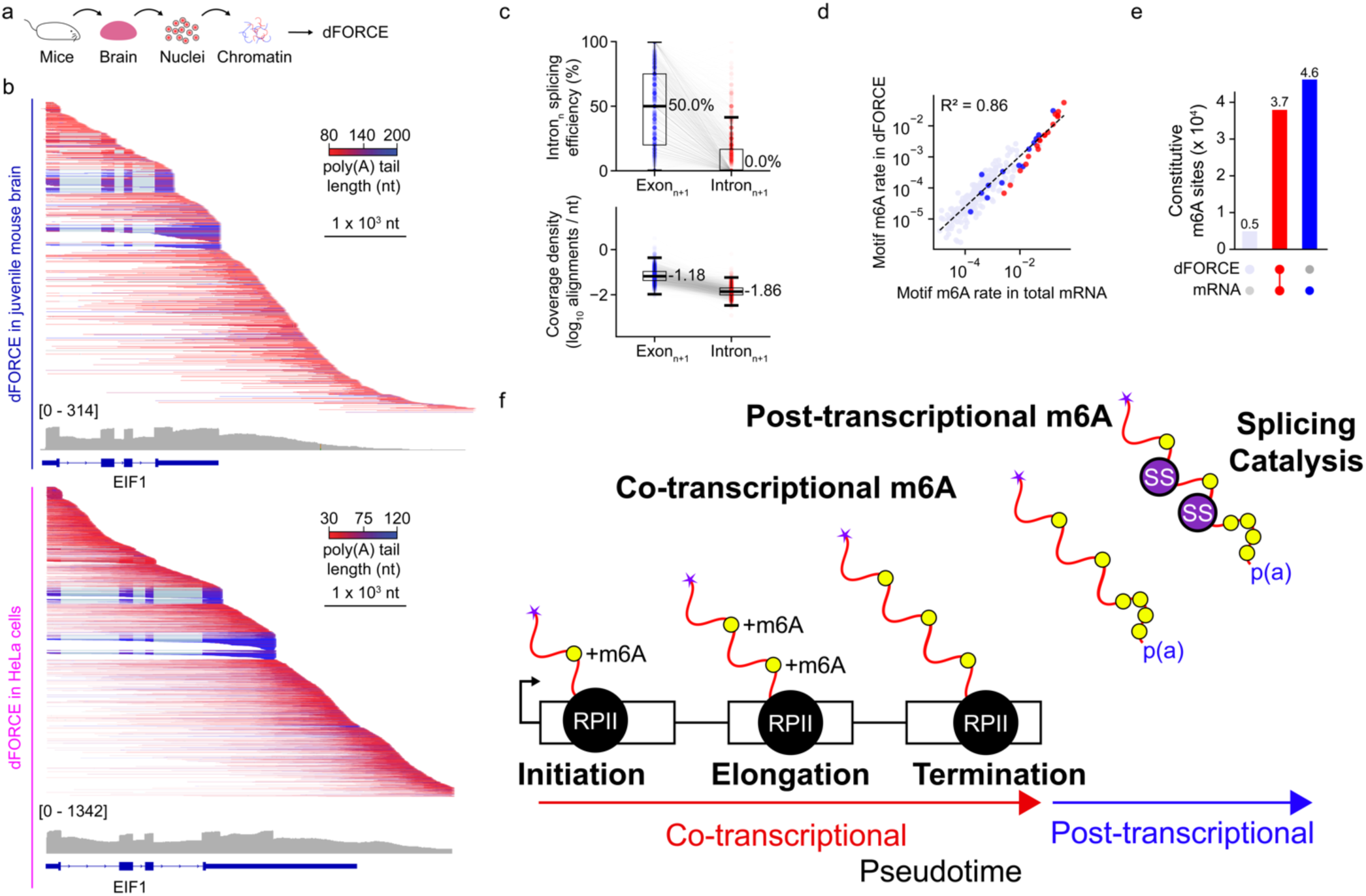
dFORCE maps the *in vivo* timing of mRNA synthesis. **(a)** Schematic of *in vivo* dFORCE in mouse brain. **(b)** dFORCE alignments at the orthologous *EIF1* locus in juvenile mouse brains (top) and HeLa cells (bottom). Displayed alignments correspond to GRCm39 region approx. chr11:100,210,500-100,215,998 and GRCh38: region approx. chr17:41,688,300-41,693,796. Alignments shown are on the plus strand, coloured by poly(A) tail length and sorted by aligned 3’ end position. **(c)** (Top) Proportion of intronn splicing seen in reads ending in the downstream exon and intron, across 32,027 intron-exon-intron trios corresponding to 6,968 mouse genes. (Bottom) Total density of 3’ end alignment terminating in same set of exons and introns. The density was calculated as the number of alignment 3’ ends divided by the feature width. **(d)** 5mer motif level m6A methylation rate in total mRNA (x-axis) and dFORCE pre-mRNA (y-axis) reads. Density calculations and colour scale correspond to panel (a). **(e)** UpSet plot representing the distribution of m6A sites in dFORCE only (light blue), total mRNA only (dark blue), or both (red), for 8.8 x 104 constitutive m6A sites (methylated at ≥ 30% stoichiometry in either dFORCE or total mRNA) which were tested in both fractions. **(f)** A schematic model representing the timing RNA polymerase II transcription, m6A deposition, 3’ end processing and splicing catalysis during mRNA synthesis of mammalian genes less than 10 kb in length. RNA polymerase II (black circle) transcribes pre-mRNA (red squiggle) in the presence of co-transcriptional m6A deposition (yellow circle) but typically in the absence of mature spliceosomes and splicing catalysis (purple circle). The pre-mRNA is fully unspliced at the time of cleavage and polyadenylation. 5’ end of pre-mRNA is indicated with a purple star. Only steps directly measured by dFORCE are represented in this plot; the specific timing of 5’ capping and recruitment of U1/U2 are excluded from this model. Pseudotime and co-transcriptional and post-transcriptional stages of mRNA synthesis are indicated at the bottom of the plot.

We next examined the kinetics of splicing during *in vivo* brain transcription. Mirroring observations in cell culture, splicing catalysis was acutely absent from pre-mRNA^e^ describing transcription through introns (Fig. 6c). The abundance of spliced pre-mRNA^e^ in the mouse brain correlated with excess alignment density in exons and was discontinuous with the extent of splicing in next introns (R^2^ = 0.90; Fig. S7e-f). Over half of spliced pre-mRNA^e^ directly corresponded to catalytic intermediates (CI*), collectively accounting for 5.5% of all pre-mRNA^e^, and the abundance of spliced pre-mRNA^e^ closely correlated with alignments ending at splice donors (*r* = 0.76), reinforcing our finding in cells that most apparent evidence for co-transcriptional splicing in fact represented CI* rather than elongating transcripts (Fig. S7g-j).

We then assessed splicing during late transcription, finding that pre-mRNA^e^ remained over 90% unspliced across 10,000 genes. Splicing was predominantly deferred until after cleavage and polyadenylation, at which point we detected abundant partially and fully spliced transcript (Fig. S8a-c). Supporting this, we identified 2,388 genes where all introns were unspliced during late transcription and 1,403 where all introns were unspliced after polyadenylation, confirming our finding in cells that splicing was often entirely post-transcriptional across thousands of genes (Fig. S8d). Finally, we estimated the extent of upstream splicing during deep intronic transcription, finding that pre-mRNA^e^ remained > 90% unspliced up to maximum observed transcriptional distances of 11 x 10^3^ nt beyond 3’ splice sites (Fig. S8f), corroborating that our previously identified delay in splicing catalysis is also present during endogenous transcription in tissues.

Assessing m6A methylation, we mapped 1.7- and 2.0 x 10^5^ modified sites across pre-mRNA and total mRNA fractions, with both sets displaying METTL3 hallmarks (S9a-d). We identified 32-fold m6A suppression on pre-mRNA introns and confirmed efficient cytoplasmic display of exonic m6A methylation (Fig. 6d and Fig. S9b-c). Similar to cell-line data, catalytic intermediates were m6A hypermethylated compared to elongating pre- mRNA^e^, with over 3000 genes showing CI* methylation (Fig. S9g). Further, median m6A stoichiometries approached 75% of cytoplasmic levels on unspliced pre-mRNA^a^ and matched or exceeded total mRNA levels on partially spliced and fully spliced pre-mRNA^a^, confirming our finding in cells that much of m6A deposition occurred prior to the commencement of splicing catalysis (Fig. S9h-i). Notably, a subset of sites displayed increased m6A on spliced pre-mRNA compared to total mRNA, suggesting a possible role for mRNA m6A demethylation^42^ during *in vivo* neuronal mRNA metabolism.

To illustrate the capabilities of *in vivo* dFORCE, we highlighted the multimodal timing of mRNA synthesis at *Npas4*, a transcriptional regulator linked to human neuropsychiatric disorders^43^. *Npas4* presented a nearly uniform pre-mRNA^e^ coverage across the gene body, consistent with ongoing transcription and nascent 3′ end formation (Fig. S10a-b). Although we could not fully resolve cleavage kinetics, termination was around 50% efficient by 1.4 kb downstream of the PAS (Fig. S10c). Spliced pre-mRNA^e^ were detected ending in all exons, but no intron-ending alignments displayed splice junctions, indicating that *Npas4* pre-mRNA remained unspliced during transcription within our sequencing lengths (Fig. S10d-e). Ranking individual pre-mRNA alignments by their m6A content suggested that transcripts with higher m6A levels were more likely to exhibit splicing, transcriptional, and polyadenylation (Fig. S10f). Taken together, our results underscore how dFORCE resolves the precise timing of transcription, splicing, and modification within an essential neuronal gene, illuminating a distinct order of events, where m6A deposition occurs early and splicing catalysis is delayed until the post-transcriptional maturation of pre-mRNA.

## Discussion

We introduce dFORCE, a direct pre-mRNA sequencing technique that illuminates the multimodal trajectory of endogenous mRNA synthesis, from RNA polymerase II transcription through multi-step splicing, m6A deposition, and 3′ end formation. By coupling large-scale RNAPII IP with direct RNA sequencing, dFORCE circumvents the possible effects of metabolic labelling, amplification, or partial fragmentation on pre-mRNA sequencing. We summarise our findings in a model where, within the genomic context we could observe, at least 20-30 thousand human and mouse introns remain largely unspliced throughout transcription, 3’ end formation and m6A methylation, with splicing catalysis taking place at a slower rate than previously proposed on polyadenylated and m6A methylated pre-mRNA substrates (Fig. 6f).

dFORCE separates RNAPII-associated non-polyadenylated and polyadenylated fractions, assessing co-transcriptional processing on genuinely elongating pre-mRNA transcripts. This allowed us to simultaneously detect 3’ and 5’ cleavage events on the same molecules. We provide key validation and extension of recursive polyadenylation^37^ which appears to play a widespread role in shaping mRNA 3’ end topology, and preliminary single-molecule estimates of the dynamics of RNAPII 3’ cleavage and termination. Our observation of widespread recursive polyadenylation implies that a large part of cleavage and polyadenylation occurs post-transcriptionally on already polyadenylated substrates, suggesting an extended window of opportunity for mRNA 3’ end selection.

Our findings indicate the need for a revaluation of the rapid co-transcriptional splicing model, which posits that splicing catalysis occurs in close proximity to elongating RNAPII^1,2,9^. Our results suggest that the catalytic intermediates of splicing, whose 3’ ends are exposed before exon joining, confound the sites of elongating RNAPII within exons, thereby creating an apparent signal of co-transcriptional splicing. Supporting this interpretation, reads ending in an intron are consistently unspliced or less frequently spliced compared to reads ending in the upstream exon, and the abundance of splicing catalytic intermediates directly correlates with the frequency of observed co-transcriptional splicing genome wide.

Our analysis on elongating pre-mRNA demonstrates that introns remain predominantly unspliced to a lower bound of 10,000 nt behind transcription. While this lower bound is imposed by read length limitations, future sequencing technologies will likely dissect the exact distances behind elongating polymerases where upstream splicing becomes predominant. Despite these limitations, we find that close to 10,000 genes in human cell lines and mouse tissues remain approximately 90% unspliced during late transcription within the context lengths provided by dFORCE. Further, we identified widespread occurrence of completely unspliced pre-mRNA both during late transcription and after polyadenylation, indicating that splicing catalysis could be entirely post-transcriptional for over 2000 multi-exon genes in human cells and mouse tissues. However, it remains unclear whether this delay corresponds to a slow spliceosome assembly or to the shuttling of unspliced pre-mRNA to nuclear speckles^44,45^.

Our model is consistent with previous studies describing a stimulatory role of poly(A) tails in driving last intron splicing^46,47^, which might extend to upstream introns. It is also consistent with recent genome-wide estimates of typical intron half-lives beyond 30 minutes, and typical chromatin residence time of pre-mRNA and mRNA in the range of 1 hour^48^. On the other hand, while the combination of 5’ splice sites and poly(A) tails marks pre-mRNA for nuclear degradation^49^, how unspliced pre-mRNA is selected for further processing or nuclear degradation remains unclear.

Importantly, our results remain consistent with previous estimates of splicing delay measured by pharmacological inhibition and release of transcription (5 to 10 min)^50^ and microscopy-based inspection of intron removal events (∼8 min)^51^. These timing delays correspond to 10 to 40 kb of transcription distance, given a transcription speed of 2 to 4kb/min^50–52^. While co-transcriptional splicing remains plausible in long introns or far away from the elongating RNAPII and may have regulatory functions^50,51,53,54^, our study shows that it may not be a mandatory or rapid process for thousands of human and mouse genes.

Importantly, our results do not exclude co-transcriptional spliceosome recruitment^4,8^. Single-molecule imaging indicates that splicing often occurs post-transcriptionally, but near the site of transcription^55^. If these post-transcriptionally partially spliced pre-mRNAs remain bound or in close proximity to RNAPII, this could explain why 15-20% of genic dFORCE alignments corresponded to endogenously polyadenylated pre-mRNAs and their catalytic intermediates. Sequestration of RNAPII with unprocessed pre-mRNA would represent an elegant method to buffer transcriptional activity against post-transcriptional processing flux, but little is known about RNAPII localisation after termination and whether it remains associated with polyadenylated but unspliced pre-mRNA.

We show that m6A accumulates to near cytoplasmic levels on unspliced and partially spliced RNAPII-associated polyadenylated pre-mRNA, as well as splicing catalytic intermediates, indicating that m6A deposition substantially precedes splicing catalysis. Critically, m6A is suppressed on exon boundaries across all fractions, including unspliced pre-mRNA, indicating that the EJC is not responsible for establishing m6A silencing at exon boundaries, as previously suggested^23–25^. It is likely that other complexes, such as snRNPs already bound to pre-mRNA before splicing catalysis, instead exclude m6A at these regions^28^. Still, our results do not exclude a role for the EJC in maintaining m6A suppression after splicing catalysis. Our findings are consistent with prior reports that demonstrate exon-specific m6A methylation on unspliced, intron-harbouring mRNA^56,57^, and m6A silencing not always associated with exon boundaries^23,58^.

dFORCE offers an integrated view of transcription-coupled RNA processing, including cleavage, polyadenylation, termination, splicing, and RNA modification including but not limited to m6A methylation, on a genome-wide scale. Our findings prompt a re-evaluation of the canonical models of splicing and RNA modification kinetics during mRNA synthesis. By leveraging single-molecule resolution, our approach sets the stage for the future mechanistic dissection of how splicing is deferred and how RNA modification occurs early on, paving the way for new insights into nuclear architecture, mRNA regulation, and transcriptomic diversification. We anticipate that dFORCE will spur fresh strategies for dissecting the molecular choreography of splicing and epitranscriptomic regulation during *in vivo* mRNA synthesis.

## Acknowledgements

We thank Prof. Hiroshi Kimura for his provision of CMA601 antibodies.

## Data availability

All datasets used in this study are publicly available. Nano-COP datasets^10^ based on Nanopore direct RNA sequencing from K562 cells were retrieved from SRA (https://www.ncbi.nlm.nih.gov/bioproject/PRJNA507789) (accessions SRR10097603, SRR15529845). HeLa POINT-nano datasets^1^ were retrieved from SRA (https://www.ncbi.nlm.nih.gov/bioproject/PRJNA668366) (accessions SRR12802908, SRR12802909, SRR12802910). Sequencing data produced in this work has been deposited to SRA (https://www.ncbi.nlm.nih.gov/bioproject/PRJNA1256087).

## Code availability

Complete code to reproduce our results is available at: https://github.com/comprna/dFORCE.

## Methods

### Definitions

- **pre-mRNA:** Precursor RNA molecules that are partially transcribed and/or partially processed compared to mature mRNA.
- **pre-mRNA^e^**: Partially transcribed molecules actively engaged with RNAPII and lacking endogenous poly(A) tails.
- **pre-mRNA^a^**: Fully transcribed, endogenously polyadenylated pre-mRNA molecules.
- **Catalytic intermediates (CI*)**: Non-polyadenylated splicing products (ending within 1 nt of an active 5′ splice site).
- **Fragmented spliced RNA**: Non-polyadenylated RNAs that may arise from endonucleolytic cleavage of spliced mRNA
- **Context length**: The confidence interval made for a claim about the range at which at pre-mRNA is unspliced in a current class (e.g. pre-mRNA^e^ ending in the last intron of a gene), based on the maximum observed alignment length corresponding to that class
- **Co-transcriptional splicing:** Splicing catalysis that occurs during RNA polymerase II transcription before cleavage and polyadenylation.

### Mammalian cell culture

HeLa cells (human cervical cancer) and K562 cells (chronic myeloid leukemia) were originally obtained from ATTC. Cells were grown in DMEM medium (Gibco) supplemented with 10% FBS and 1% antibiotic-antimycotic solution (Sigma-Aldrich) and passaged when 70–90% confluent. For the METTL3 inhibition, HeLa cells were cultured with 10 μM final concentration of METTL3 inhibitor (STM2457, MedChemExpress) for 2 hours.

### Mouse husbandry

All experimental procedures were conducted using C57BL/6J adult male mice (4-5 weeks old), group-housed in controlled environmental conditions at the Australian Phenomics Facility (APF), Australian National University. The mice were temporarily housed in a holding room (12-12 h reverse light/dark cycle, 22°C ± 2°C) with food and water available *ad libitum*. All experimental protocols were approved by the Animal Experimentation and Ethics Committee of the Australian National University (A/2023/304) and conducted in accordance with the National Health and Medical Research Council’s (NHMRC) Australian Code for the Care and Use of Animals for Scientific Purposes and the Australian Capital Territory Animal Welfare Act 1992.

### *In vitro* RNA polymerase II immunoprecipitation

RNA polymerase II immunoprecipitation was performed as previously described in the POINT protocol with the following modifications^1^. 0.2 ∼ 1.0 x 10^8^ exponentially growing HeLa and K562 cells were harvested and lysed in 4 ml of HLBN buffer (10 mM Tris-HCl, pH 7.5, 10 mM NaCl, 2.5 mM MgCl_2_ and 0.5% v/v NP-40, supplemented by cOmplete EDTA-free protease inhibitor cocktail, Sigma) on ice for 5 minutes. Crude cell lysates were underlaid with 1 ml of HLBNS buffer (HLBN buffer + 10% w/v Sucrose) and spun at 300 g for 5 minutes to collect nuclei. Nuclei were washed by gentle resuspension in 1 ml NUN1 buffer (20 mM Tris-HCl, pH 7.9, 75 mM NaCl, 0.5 mM EDTA and 50% v/v glycerol), overlay of the resuspension on 1 ml of clean NUN1 buffer, and centrifugation at 3000 g for 2 minutes. This wash step was repeated twice. Purified nuclei were finally resuspended in 400 µL NUN1 buffer. At this step, for experiments in K562 cells, we spiked-in 1 μg of *Drosophila* S2 cell poly(A) RNA. Nuclear lysis was performed by the gentle addition of 4 ml NUN2E buffer (20 mM Hepes-KOH, pH 7.6, 300 mM NaCl, 0.2 mM EDTA, 1% v/v NP-40, 1M Urea and 3% v/v Empigen, Sigma). Samples were slowly inverted twice to mix the solutions, and reactions were gently rocked on ice for 5 minutes, during which chromatin formed large wispy strands which slowly condensed into a gel-like droplet. Delicate handling during chromatin gel formation was essential to prevent chromatin aggregation, which would cause chromatin to precipitate out of solution during DNase digestion.

After formation, the chromatin gel was fished from the NUN2E using a P1000 pipette. The chromatin gel was immersed in 45 ml of ice-cold PBS and washed by gentle inversion and incubation on ice for 1 minute. This wash was repeated twice, and then the chromatin gel was transferred to 45 ml of high-salt DNase buffer (10 mM Tris-HCl pH 7.5, 400 mM NaCl, 100 mM MnCl_2_) for buffer equilibration. Buffer equilibration was performed by gentle inversion and incubation on ice for 2 minutes. Finally, the chromatin gel was added to 200 µL of turbo DNase reaction (10 mM Tris-HCl pH 7.5, 400 mM NaCl, 100 mM MnCl_2_, 0.02 U/μL SUPERase·In and 0.2 U/μL Turbo DNase, Thermo Fisher) and digested for 15 minutes at 37 °C while shaking at 1200 rpm. The chromatin dissolved completely after 7 minutes. Reactions were quenched by the addition of 8 μl of 0.5 M EDTA and brief pulse vortexing. Reactions were clarified by centrifugation at 16,000 g for 3 minutes, before collection of soluble chromatin contained in the supernatant. Transcription complexes were immunoprecipitated using 10 μg of CMA601 RNAPII CTD-targeting antibodies (RRID:AB_2827956) bound to 200 μl of Dynabeads M280 anti-mouse IgG (Invitrogen; 11201D) in 1.5 ml of NET2E buffer for 1 – 2 hours at 4 °C on a hula mixer. 10 μg of anti-Drosophila Aubergine antibodies (mouse monoclonal 8A3-D7, gift from Julius Brennecke) were used as a control for western blotting. Beads were washed 6 times with 1 ml of 4 °C NET2E buffer.

Following the wash, beads were gently resuspended in 300 μl Tri Reagent (Sigma) and held on a HulaMixer rotating tube mixer (Invitrogen) for 5 minutes at room temperature to release RNA and protein. Beads were captured and the supernatant was used to extract RNA using the Direct-Zol RNA MicroPrep kit (Zymo Research), following manufacturer’s instructions, including the optional on-column DNase digest. RNA was eluted in 10 μL water and immediately polyadenylated with 1 μL of bacterial poly(A) polymerase (5 units; New England Biolabs #M0276S) in a 20 μL reaction. Polyadenylation was conducted for 2 minutes at 37 °C, and brief timing of the polyadenylation was essential to distinguish between *in vitro* and *in vivo* poly(A) tails on the basis of length. Polyadenylation reactions were quenched by the addition of 5 μL of 50 mM EDTA, and RNA was purified by the addition of 45 μL of AMPure XP SPRI beads (Beckman Coulter), following manufacturer’s instructions. Polyadenylated RNA was eluted in 10 μL water and taken forward for library preparation.

### Preparation of mouse brain for RNA polymerase II immunoprecipitation

Mice were deeply anesthetised with isoflurane before euthanasia via decapitation and immediately the brain was dissected without fixation. To prevent RNA degradation and ensure protein stability, brains were rapidly extracted within 2 minutes post-euthanasia and immediately immersed in ice-cold phosphate-buffered saline (PBS; pH 7.4). Brains were transferred into 8 ml of HLBN lysis buffer and homogenised by 30 strokes in a tight 15 ml douncer, followed by incubation on ice for 10 minutes with intermittent douncing. The crude brain lysate was then successively filtered through 500 μm (Sigma-Aldrich; CLS3477) and 70 μm cell strainers (Falcon; 352350) to remove large debris. The strained lysate was overlaid on 1 ml of HLBN+S and centrifuged at 300 g for 5 minutes at 4 °C to collect the crude nuclear fraction. From this step, Pol II IP was processed identically to the *in vitro* experiments.

### Western blotting

Cytoplasmic, soluble chromatin and the immunoprecipitation fractions from the POINT experiments were loaded on NuPage 4–12% Bis-Tris Protein Gels (Invitrogen) followed by transfer onto PVDF membrane. The membrane was blocked in Odyssey Blocking Buffer (for IR-Dye detection; LI-COR 927-40000) and probed with primary antibodies: CMA601 RNAPII CTD-targeting antibodies (1:5,000 dilution), anti-Histone H3 antibodies (Ab1791, Abcam, 1:1,000 dilution) and anti-GAPDH antibodies (Ab125247, Abcam, 1:1,000 dilution). The membranes were then probed with a secondary antibody, either anti-mouse-IR-Dye800 (1:10,000, 926-32210; LI-COR) or anti-rabbit-IRDye680 (1:10,000, 925-68071; LI-COR), and imaged using the Odyssey CLx Imaging System (LI-COR).

### Total RNA isolation from HeLa, K562 cells and mouse brain

Exponentially growing HeLa and K562 cells were washed in ice-cold PBS and collected by centrifugation at 300 g for 5 minutes. Cells were lysed by the addition of Tri Reagent (Sigma-Aldrich; T9424), followed by hula mixing for 5 minutes at room temperature. Tri reagent was clarified by centrifugation at 10,000 g for 1 minute at room temperature. RNA was extracted from the soluble fraction using Direct-Zol RNA MicroPrep kit following manufacturer’s instructions, including the optional DNase step. Total RNA was eluted in 10 μl of water. 1 μg of total RNA was taken forward for sequencing without *in vitro* polyadenylation or poly(A) selection.

Mouse brain total cell lysates were collected after homogenisation with the HLBN lysis buffer. ∼50 μl of lysates were mixed with 300 μl of Tri Reagent, followed by hula mixing for 5 minutes at room temperature. Total RNA was purified using Direct-Zol RNA MicroPrep kit.

### Nanopore direct RNA library preparation and sequencing

Direct RNA sequencing was performed on a PromethION P2 solo sequencer using PRO004RA flow cells in two software configurations: *long-fragment* (minKNOW 23.11.7) and *all-fragment* (minKNOW 24.06.10). MinKNOW 23.11.7 software only sequenced RNA strands of greater than 200 nt in length, whereas MinKNOW version 24.06.10 sequenced RNA as short as 50 nt. RNA libraries were prepared using the direct RNA sequencing kit (ONT; SQK-RNA004) following manufacturer’s instructions, with the omission of the RNA control standard and substitution of Induro reverse transcriptase (NEB; M0681) in place of the recommended reverse transcriptase. RNA was reverse transcribed for 15 minutes at 60 °C and immediately cooled to 4 °C, omitting the denaturation of the reverse transcriptase to preserve RNA integrity. 1 μl of SUPERase·In was added for the reverse transcription adaptor ligation step.

For POINT experiments, 200 ng – 1 µg of *in vitro* polyadenylated RNA was used for library preparation, depending on experiment yield. For total RNA experiments, 1 µg of total RNA was used for library preparation. Sequencing was performed until sufficient data yield was reached, and flow-cells were washed using the ONT flow cell wash kit (ONT; EXP-WSH004) with the addition of 1 μl RNase I (ThermoFisher; EN0601). Flow cells were reloaded immediately or stored at 4 °C following manufacturer’s instructions.

### Reference genomes

Reference genome assemblies and gene structure annotations were retrieved from Ensembl^36^ (release 113) for all analysed species. For HeLa experiments, genome alignment was performed against the GRCh38 unmasked primary assembly. For K562 experiments, which were spiked-in with S2 poly(A)-selected RNA, alignment was jointly performed against GRCh38 unmasked primary assembly and the *Drosophila* BDGP6.46 unmasked top-level assembly. For mouse *in vivo* dFORCE experiments, alignment was performed against GRCm39 unmasked primary assembly.

### Modification-resolved basecalling and genomic alignment

Basecalling and alignment were performed on compute nodes equipped with Tesla Volta V100-SXM2-32GB GPUs. Basecalling was performed using Dorado v0.8.0 software and rna004_130bps_sup@v5.1.0 basecalling models, enabling detection of poly(A) tail lengths and m6A, m5C, inosine, and pseudouridine modifications with command dorado basecaller sup,pseU,inosine_m6A, --estimate-poly-a -r -b 416 -c 9216’. Following basecalling, RNA sequences were extracted from the unaligned BAM output using samtools^59^ v1.18 with command ‘samtools fastq -T “*”’. Spliced alignment was performed against reference genomes using minimap2 v2.27 with parameters ‘minimap2 -a -x splice -y’. Unmapped reads, secondary- and chimeric alignments were removed using ‘samtools view -F 2308’.

### Assignment of alignments to genes and 3’ end features

We assigned dFORCE sequencing alignments to genes based on the overlap between alignments, and filtered gene structures. We intersected aligned regions of genes (indicated by the ‘M’ cigar operator) against the transcript coordinates in a stranded manner using bedtools^60^ v2.29 with command ‘bedtools intersect - wo -s’. Intersection was performed against gene regions, exon regions and 5 kb downstream of gene regions. Alignments were preferentially assigned to the genes with which they had best exonic overlap, intronic overlaps, or downstream-of-gene overlap, in that order. Alignments which didn’t overlap any of these features were classified as intergenic. Following assignment of reads to genes, the last 6 aligned nucleotides for each read were compared to the gene structure and the entire alignment was assigned to the corresponding exon, intron, or downstream of gene feature associated with that gene, requiring a minimum overlap of 4 nt with an exon, intron or DOG feature. This assignment process was done in 3-fold, once against entire Ensembl v113 annotations (for biotype plots), then against first-pass and second-pass refined annotations (see below).

### Separation of co-transcriptional and post-transcriptional read fractions in HeLa and K562 cells

To distinguish between endogenously polyadenylated and non-polyadenylated reads, we normalised the poly(A) tail length of each pre-mRNA alignment to a background distribution derived from ribosomal RNA and small nuclear RNA alignments sequenced within the same library. These control RNAs are known not to be endogenously polyadenylated. For each library, we calculated the mean and standard deviation of poly(A) tail length for the control non-polyadenylated reads and calculated the poly(A) tail length z-score for each protein-coding (pre-mRNA) read against this distribution, for pre-mRNA alignments of at least 200 nt in length. A Z-score cutoff of ≤ 1.8 was used to classify reads as non-endogenously polyadenylated (labelled pre-mRNA^e^) and were thus considered to be co-transcriptional reads. A poly(A) Z-score of cutoff ≥ 3.6 was classified as endogenous polyadenylation, and reads were labelled as pre-mRNA^a^. Protein-coding alignments less than 200 nt in length, or with poly(A) Z-scores between 1.8 and 3.6 were excluded from classification and downstream analysis.

### Separation of co-transcriptional and post-transcriptional read fractions in mouse brain

Mouse pre-mRNA alignments (mapped to the protein-coding genes or their downstream-of-gene regions) were normalised to poly(A) tail length distributions from alignments assigned to snRNA, mtRNA, and rRNA genes. Owing to longer in vitro poly(A) tails added during in vivo dFORCE, we adjusted poly(A) z-score threshold to maintain a good separation between the fractions (Fig. S7b). Z score thresholds of up to 1.8 were classified as pre-mRNAe and of at least 4.2 were classified as pre-mRNA^a^. Minimum pre-mRNA alignment length was set to 100 nt.

### Visualisation of RNA poly(A) tail length via genome browser

Poly(A) tail lengths were captured during sequencing and quantified during basecalling using the Dorado flag ‘--estimate-poly-a’, producing a ‘pt’ BAM flag. To visualise poly(A) tail lengths on the genome browser, we extracted poly(A) tail lengths measurements from basecalling data and used it to annotate BAM files with a colour YC tag corresponding to the length of the tail. Colour tags were visualised using IGV genome browser.

### Annotation curation and isoform refinement

To classify pre-mRNA alignments as spanning intronic or exonic regions, we compared alignments to a single-representative isoform for each protein-coding gene. To select these representative transcript models, we used a multi-step filtering process using both total mRNA sequencing data and dFORCE pre-mRNA^a^ data.

1. Preprocessing Ensembl annotations

Starting with transcript models annotated in Ensembl v113, we filtered for transcript models annotated by Ensembl, HAVANA or miRbase, and further, RefSeq (for mouse analysis only). Using this filter, we generated a smaller GTF file containing only transcripts annotated from these sources.

2.First-pass classification and GTF filtering

To refine the annotation further, we computed the support for each transcript based on total RNA genome alignments. To select dominant isoforms for genes with reads, transcripts were ranked by a combined score of end support and splice support. The highest-scoring isoform was selected. For genes without reads, the longest isoform was kept.

3. Poly(A) site refinement

We next used dFORCE pre-mRNA^a^ alignments with at least 200 nt of aligned bases to perform simple poly(A) site clustering across 30 nt sliding windows of the genome. The most downstream cluster for each gene with at least 10 supporting pre-mRNA^a^ alignments was selected as the dominant poly(A) site. Using these clusters, we extended or truncated previous isoform selections to use dominant poly(A) sites as the transcript end site.

Together, these steps yielded a single refined transcript for every expressed protein coding genes, referred to as the ‘second pass index’. All subsequent analyses were performed against this second pass index. All analyses except initial biotype assignments were done using this refined ‘second-pass’ annotation.

### Clustering and selection of consensus polyA sites (PAS) from dFORCE data

We identified pre-mRNA 3’ end poly(A) site clusters (PAS) for protein-coding genes by clustering the terminal 3’ aligned coordinates of post-transcriptional (pre-mRNA^a^) reads, after they had been assigned to genes. For each group of 3’ end coordinates per gene, clusters were generated using DBSCAN with eps=5, metric=‘euclidean’, and a minimum clusters size of at least 2 alignments and at least 1% of all pre-mRNA^a^ identified per gene. Cluster positions, supporting reads, and distances to annotated poly(A) sites were recorded in BED format. Motif discovery was performed using MEME v5.5.7^61^, analysing the 40 nt sequences centered on the middle coordinates of PAS clusters. MEME command ‘meme all_clusters.fa -dna -oc . -nostatus -time 14400 -mod zoops -nmotifs 10 -minw 6 -maxw 6 -objfun classic - markov_order 0’ was used.

### Validation of dFORCE poly(A) sites against total RNA and poly(A) site annotations

PAS coordinates were 3-fold validated against poly(A) sites seen in our own total RNA sequencing, and via comparison to poly(A) sites annotated in Ensembl v113 and PolyASite Atlas 2.0^35^. For comparison to total RNA, we required at least 1 total mRNA alignment (sequenced without *in vitro* polyadenylation) to present a 3’ end within ±2 nt of the PAS midpoint. For Ensembl and PolyASite annotations, we required an annotated PAS to lie within ±20 nt of the annotated poly(A) site for it to be considered to be supported.

### Analysis of intronic poly(A) site clusters

To identify intronic poly(A) sites, we first assigned all PAS to subgenic regions, using gene structures generated in the second-pass transcript structures. For PAS sites spanning the boundaries of exons, they were preferentially assigned to exonic regions before assignment to either intronic or DOP regions. Metagene coordinates were calculated relative to second-pass gene structures. Distances between intronic poly(A) sites and CDS start positions were calculated using the annotated CDS start in Ensembl for transcript selected during second-pass annotation filtering. To study the association of cryptic splicing with PAS, we calculated the incidence of cryptic splicing (the display of splice junctions not annotated in any transcript in Ensembl v113) for pre-mRNA^a^ alignments corresponding to identified PAS. If any pre-mRNA^a^ ending at a given poly(A) site cluster presented a splicing event, the cluster was associated with spliced reads. If any pre-mRNA^a^ ending at a PAS presented a not-in-annotation splicing event (i.e., an event not annotated in any Ensembl v113 transcript), the PAS was considered to present novel splicing events.

### Detection of recursive polyadenylation

To identify recursive polyadenylation, we screened for the presence of bridging and spanning pre-mRNA^a^ alignments (Fig. 2d). To identify bridging alignments, we screened for pre-mRNA^a^ alignments whose 3’ ends ended within a window of *k* nucleotides around a downstream PAS, and whose 5’ end lay within a window of *k* nucleotides of an upstream 5’ PAS. *K* was set to 20 nt to allow for slight positional variation of alignment end coordinates around each PAS. Only pairs of poly(A) sites bridged by at least 2 pre-mRNA^a^ alignment were considered as putative bridging alignments indicating recursive polyadenylation. To identify spanning pre- mRNA^a^ alignments, we identified pre-mRNA^a^ alignments which ended at the downstream PAS of a bridged PAS pair, and had a 5’ end within the -40 to -20 region upstream of the ‘start’ coordinate of an upstream PAS. These alignments were polyadenylated at the downstream PAS yet spanned through the upstream PAS, potentially representing intermediates of the recursive polyadenylation process. For functional GO enrichment, genes presenting at least 1 bridged PAS pair were compared to the set of all tested genes. goatools^62^ was used to assess for function enrichment using the Homo Sapiens molecular function annotation^63^ using the 2024-01-17 GeneOntology release^64^.

### Classification of cleaved and uncleaved pre-mRNA^e^

To classify pre-mRNA^e^ alignments spanning dominant poly(A) sites as cleaved or uncleaved, we assessed the overlap of pre-mRNA^e^ with two windows around poly(A) sites. The first window was from the -40 nucleotide to the +0 nucleotide around the PAS, and the second window was from the +10 nucleotide to the +100 nucleotide of the PAS. Using previous alignment to gene classifications, we assessed for overlap with each of these windows, using bedtools command `‘bedtools intersect -a "$alignments" -b "$regions" -s -wa -wb -split -F 0.5 -bed’`. A minimum 50% overlap of the window was required. Alignments were modelled as the downstream products of cleavage if spanning only window 2, or uncleaved if spanning both windows. Alignments spanning only the 5’ window or neither region was excluded from the analysis. To visualise cleaved and uncleaved pre-mRNAe on genome browser, we added ‘YC’ colour tags to BAM alignments corresponding to their splicing classifications. Downstream products of cleavage pre-mRNA^e^ were coloured blue and uncleaved pre-mRNA^e^ spanning both regions were coloured green. All remaining pre- mRNA^e^ alignments were coloured grey. Alignments were visualised using IGV genome browser.

### Calculation of co-transcriptional 3’ cleavage distance (C_50_)

To determine the transcriptional timing of pre-mRNA 3’ cleavage, we first determined the distance *x* between pre-mRNA^e^ 3’ ends and the upstream poly(A) site and determined the *p_cleavage_* in each bin as 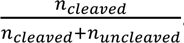. We restricted analysis to pre-mRNA^e^ where the 3’ ends within [0, +4000] nt of the PAS and retained only categories indicative of cleavage (described above), modelling the probability of cleavage *p_cleavage_* as a function of distance transcribed past the PAS, *x*, as:

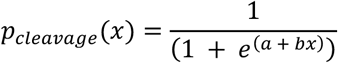

We initialised *b* as 0.001 and *a* as −*b* * *x_median_* based on the distribution of observed offsets and estimated these parameters via nonlinear least-squares optimisation with the Levenberg–Marquardt algorithm, via the scipy^65^ `‘scipy.optimize.curve_fit’` function. From this estimate, we calculated the *C*_50_ distance as 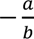, representing the transcriptional distance where 50% of elongating pre-mRNA^a^ had been cleaved at the 5’ end.

*C*_50_ estimation was only performed for genes which had at least 30 ‘uncleaved’ or ‘downstream product of cleavage’ alignments.

If the fitted *C*_50_ distance lay beyond the observed ranges of *x* values, we estimated at lower bound on the true *C*_50_ using the 99^th^ percentile of observed *x* distances. If a fit failed to converge, we reported no *C*_50_ for that gene.

### Calculation of termination distances (T_50_)

From the same data used to estimate cleavage, we also estimated transcription termination behaviour downstream of the PAS, based on the decaying density of pre-mRNA^e^ 3’ ends. We restricted *x* to the range [+50, +4000] nt to exclude signals very close to the PAS and to avoid spurious signals further downstream. We binned the data in 200 nt intervals and computed the density of pre-mRNA^e^ 3′ ends within each bin. We normalized each bin’s coverage by the maximum coverage across all bins to obtain a coverage fraction. The local maximum bin was identified, and the “peak coverage” *cov_maximum_* was defined as the highest normalised coverage; the median coverage up to that peak bin (*cov_medium_*) was then computed to anchor subsequent estimates. From these parameters, we fit a logistic decay function to pre-mRNA^e^ 3’ coverage downstream of the peak coverage:

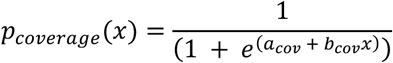

Only bins where the coverage reached at least 5% of *cov_maximum_* were considered to minimise noise in extremely low coverage regions. We identified the start of the decay regions by scanning the smoothed coverage to find the point at which it began to decrease, using a simple three-bin moving average to locate where coverage begins to decrease steadily. We initialised *b_cov_* as 0.001 and *a_cov_* as −*b_cov_* * *x_median_*. The *T*_50_ distance was then estimated as the x-value where *p_coverage_*(*x*) decreased to half of *cov_median_*. If no suitable decrease was identified or insufficient bins were above the threshold, no termination fit was performed. If the *T*_50_ solution did not fall within [+50, +3800] nt of the PAS, if fewer than six bins had pre-mRNA^e^ observations, or if the logistic fit did not converge, we reported no *T*_50_ distance for that gene.

### Summarisation of spanned exons and introns

Following previous assignments of alignments to genes, the overlap between aligned regions (represented by ‘M’ cigar operator) and exons, introns and downstream-of-gene regions of the assigned gene were calculated. As a minimum overlap threshold, only regions with at least 7 nt overlap between a given pre-mRNA alignment and an annotated genic feature were considered. From this data, we summarised the introns and exons to which a read overlapped. Additionally, the number of aligned splice junctions (cigar ‘N’ operator) were recorded.

### Identification of cryptic splicing

Given the summarised information about the set of exons and introns spanned by an alignment, along with the number of splice junctions, we identified and excluded cryptic splicing. To do so, we analysed the transition pattern between exons and introns. Alignments spliced as non-canonical compared to the annotation (e.g. contained splice junctions that were inconsistent with annotated exon-intron boundaries) were excluded. The algorithm performs an additional validation by backtracking through the read’s genomic features from its 3’ end feature, confirming that each feature was correctly preceded by its expected counterpart (i.e., exons preceded by previous intron or exon, where an upstream feature was present) according to the gene model’s architecture. Any inconsistency in this ordered progression results in a cryptic splicing classification.

### Classification of pre-mRNA^e^ upstream splicing status

Using the assignment of overlapping exons and introns per pre-mRNA alignment, we classified the splicing of all introns observed in the read. To assess the splicing of an intron, we required that the intron was fully transcribed, i.e. that the +1 nucleotide of the splice acceptor was present in the alignment. Therefore, intron-ending reads that were fully spliced at upstream introns were classified as “fully spliced”; and exon-ending reads where only a single upstream intron was spanned and was retained were classified as fully unspliced. Alignments entirely contained within an intron or only spanning a single 5’ splice site and ending in the intron were classified as ambiguous, since the intron was partially transcribed and could not be spliced. Alignments where at least one unspliced intron was observed and no exon-junctions were observed were classified as unspliced, alignments where at least 1 exon junction was observed and no unspliced, but fully transcribed introns were observed were classified as ‘fully-spliced’, and alignments where at least 1 spliced intron and at least 1 unspliced intron were observed were classified as ‘partially-spliced’.

### Classification of single intron splicing for intron-exon-intron trio analysis

For the intron-exon-intron trio splicing analysis, we separately classified the splicing status of a single intron, intron_n_, for alignments ending in either the next exon_n+1_ or the next intron_n+1_. This classification was performed similarly to above, where pre-mRNA^e^ alignments ending in exon_n+1_ or intron_n+1_ were classified as ‘unspliced’ with respect to intron_n_ if they spanned it’s 3’ terminus, and ‘spliced’ with regards to intron_n_ if they spanned exon_n_ and exon_n+1_ but not intron_n_, indicating the splicing of these intron. Pre-mRNA^e^ ending in exon_n+1_ or intron_n+1_ which did not reach the upstream intron_n_ were classified as ambiguous and excluded from the analysis.

### Calculation of pre-mRNA^e^ feature density for intron-exon-intron trio analysis

For the intron-exon-intron trio splicing analysis, we selected co-transcriptional pre-mRNA^e^ reads and calculated the sum of reads aligning to given exon, intron and downstream of gene regions, calculating density as the sum of alignments ending in a region, divided by the length of the region in the principal transcript isoform we previously refined. For downstream of gene regions, their length was initialised at 5 kb and trimmed in cases where another gene on the same strand overlapped with the DOG region.

### Measurement of intron splicing efficiency across intron-exon-intron trios

Following calculation of intron_n_ splicing status for each pre-mRNA^a^ alignment ending in a downstream exon or intron (i.e., any exon and adjacent downstream intron with rank greater than 1), and the calculation of pre- mRNA^e^ 3’ end density in each exon and intron, we summarised the splicing efficiency of each intron_n_ across adjacent intron-exon-intron trios. Several filters were used before presenting this data, namely, at least 4 pre- mRNA^e^ alignments needed to have a classified intron_n_ splicing status in both of exon_n+1_ and intron_n+1_, and that all features had a width of at least 30 in our annotation. We also excluded trios where more than 20% of alignments spliced into a 3’ poly(A) tract, rather than a conventional -AG splice acceptor, a result which occasionally occurred due to basecalling and aligning errors in the direct RNA sequencing technology.

### Reanalysis of intron-exon-intron splicing patterns from published sequencing datasets

To reassess co-transcriptional splicing discrepancies in previous reports, we downloaded and reanalysed sequencing data from studies which previously performed long-read sequencing of nascent RNA in human cell culture^1,10^. Included samples and studies are described in Table S1. Basecalled sequencing data was downloaded in FASTQ format and analysed in the same manner as our dFORCE data, without differentiating between pre-mRNA^e^ and pre-mRNA^a^, since poly(A) tail basecalling was not uniformly available from the published datasets. We compared the alignments to our custom annotation produced via dFORCE experiments in HeLa cells. For compatibility with our existing pipeline, which was designed for stranded data, only sense alignments were assigned to protein-coding genes, leading to a loss of 50% of coverage for cDNA libraries that did not use strand-specific sequencing technologies. This approach could lead to some mis-strandedness for cDNA libraries.

### Annotation of alternative 5’ splice sites that are active in pre-mRNA

In order to accurately identify splicing catalytic intermediates (defined below), we had to identify principal and alternative 5’ splice sites that were used in our samples. To do so, we extracted all active introns from our dFORCE data. We parsed ‘N’ operations from alignment cigar strings of primary dFORCE alignments. Junctions with at least 2 supporting reads were kept. The resulting set underwent a second filtering step that removed introns below a relative frequency threshold of 0.002, i.e. overlapping junctions that had fewer than that proportion of supporting reads were removed. Overlapping introns that shared more than 60% of their length were compared by their relative usage; those with a five-fold lower usage ratio were removed.

### Classification of splicing catalytic intermediate (CI*) alignments

To separate out 5’ catalytic intermediates (CI*) from other pre-mRNA^e^ classified as being engaged in transcriptional elongation, we calculated the stranded distances between pre-mRNA^e^ alignment 3’ end coordinates and the set of previously identified constitutive and alternative 5’ splice sites. Pre-mRNA^e^ alignments with 3’ ends terminating within 1 nt of an identified 5’ splice site we classified as CI* alignments.

### Calculation of gene and region-specific context lengths for splicing analysis

For each combination of region and poly(A) tail status, e.g. co-transcriptional pre-mRNA^e^ ending in the last introns, we calculated a context length for each gene. The context length was calculated as the maximum observed alignment length for reads of that gene and poly(A) tail classification ending in that window.

### Estimation of co-transcriptional splicing efficiency during deep intronic transcription

To estimate the efficiency of upstream co-transcriptional splicing catalysis and large distances relative to elongating RNA polymerase II, we screened for alignments spanning at least one fully-transcribed intron (indicated by an alignment spanning the +1 nucleotide of a splice acceptor, and at least 7 nucleotides of upstream sequence, be it the unspliced intron or the preceding exon). We then identified continuously spanned distances between transcribed 3’ splice sites and pre-mRNA^e^ 3’ end termini, indicating the absolute distance traversed by Pol II following transcription of a 3’ splice site. For each such distance, we classified the upstream 3’ splice site as either spliced or unspliced, depending on whether the region upstream of the +1 nucleotide of the splice acceptor indicated the presence or absence of splicing or not. In the case where a single pre-mRNA contained multiple adjacent unspliced introns continuous with the 3’ end of the alignment, all pairs of distances were used for the visualisation.

### Classification of m6A-methylated sites

Following basecalling for m6A modifications, RNA modification sites were assessed from primary genomic alignments, using `modkit 0.4.1 with command ‘modkit pileup –filter-threshold 0.9 --mod-threshold "a:0.99 --mod-threshold "m:0.99” --mod-threshold "17596:0.99” --mod-threshold "17802:0.99”’`. Sites were filtered for predictions where the aligned base matched the reference base. Modified sites in total RNA were defined as reference ‘A’ positions if at least 5% of reads spanning that position were m6A methylated at that site and at least 2 reads with m6A methylation were observed. For dFORCE data, we first filtered for pre-mRNA (alignments assigned to protein-coding genes) before performing the same analysis. For subsets of dFORCE alignments (e.g. unspliced pre-mRNA^e^, we filtered for read IDs previously carrying those assignments before running modkit.

### Benchmarking of m6A sites against GLORI

m6A sites previously identified by GLORI^41^ in HEK293 total mRNA were compared to those identifies in this work in HeLa total mRNA using direct RNA sequencing. Sites with at least 5 average coverage in GLORI and 5 assessed reads in HeLa total mRNA were compared.

### Visualisation of m6A sites on single pre-mRNA molecules

To view m6A sites on the genome browser, we used a combination of modkit and custom python scripts to only retain MM and ML bam tags for nucleotides that were classified as m6A positive. First, we used modkit to quantise m6A predictions on primary alignments using command ‘modkit call-mods –filter-threshold 0.9 --mod-threshold "a:0.99 --mod-threshold "m:0.99” --mod-threshold "17596:0.99’. We then filtered out 0-probability MM/ML tags and those corresponding to inosine, pseudouridine or m5C.

### Metatranscript m6A visualisation

To assign m6A sites to transcript isoforms, total mRNA reads from protein-coding genes were re-aligned to cDNA transcripts (GRCh38 or GRCm39, Ensembl release 113) using minimap2 v2.27 with parameters ‘-ax map-ont -k14 -y’. Unmapped, secondary, supplementary and negative-strand alignments were excluded using ‘samtools view -b -F 2324’. From remaining alignments, m6A sites were called as before using modkit v0.4.1, yielding m6A sites with isoform-specific coordinates. Methylation calls were annotated with distances to splice junctions and metatranscript coordinates using R2Dtool^66^ v2.0.1 using default parameters. Metatranscript and metajunction plots were produced using R2Dtool v2.0.1.

### Differential m6A analysis

Sequencing alignments were filtered based on poly(A) status and as described above. Differentially modification testing was performed using ‘modkit dmr pair –base a’. Testing was only performed on sites known to be m6A positive in total mRNA, and test results were adjusted for multiple comparisons using Benjamini/Hochberg FDR correction. Differentially modified m6A sites were defined as those where the change in m6A stoichiometry (Δm6A) was at least 10% and where FDR ≤ 0.1.

### Comparison of pre-mRNA fractions between orthologous mouse and human loci

After processing dFORCE in HeLa cells and mouse brains, we compared the proportion of pre-mRNA^e^ and pre-mRNA^a^ per gene at one-to-one orthologue gene pairs, which were retrieved from Ensembl BioMart. We selected gene pairs with at least 1000 pre-mRNA alignments in each species.

