## Supplementary Figures for "Single-molecule multimodal timing of *in vivo* mRNA synthesis"

### Figure S1

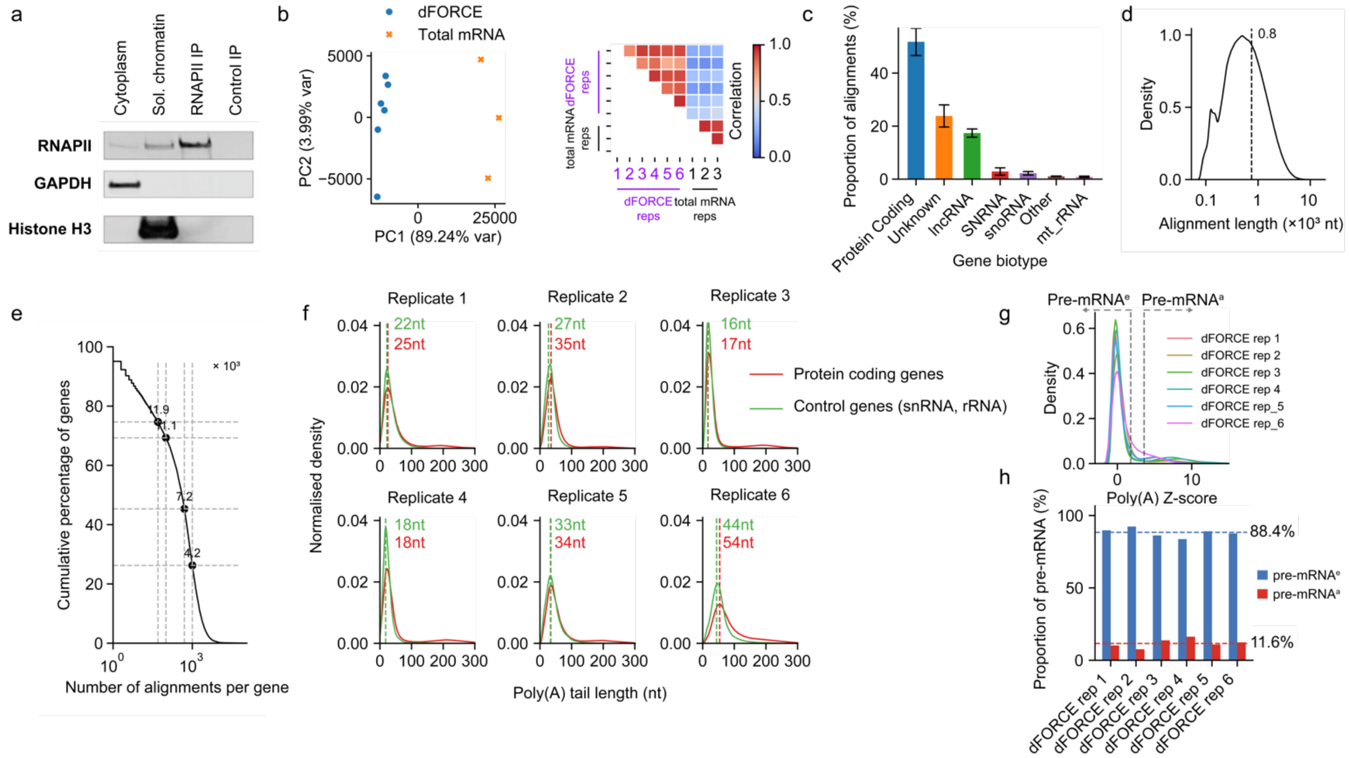


**Fig. S1. Supplemental information for HeLa dFORCE and fractionation of polyadenylated pre-mRNA. (a)** Western blotting against RNAPII CTD, GAPDH and Histone H3 antibodies across HeLa cytoplasm, soluble chromatin, RNAPII CTD IP and control antibody IP (anti-Drosophila Aubergine) fractions. 0.3%, 2.5%, and 25% of the cytoplasmic, the soluble chromatin, and the IP fractions were loaded. **(b)** PCA (left) and correlation plot (right) for TPM-normalised gene abundance in HeLa dFORCE and total mRNA sequencing for n = 11,839 genes. **(c)** Distribution of biotypes for dFORCE alignments, comparing alignments to Ensembl v113 GRCh38 annotation. Error bars represent the standard error of the mean across 6 biological replicates. **(d)** Distribution of alignment length for 1.32 x $10^7$ dFORCE alignments mapped to protein-coding genes. **(e)** Distribution of alignments per gene for 15,951 protein-coding genes measured in dFORCE. **(f)** Distributions of poly(A) tail lengths measured from reads associated with protein-coding genes (red) and non-polyadenylated control genes (green) (snRNA, rRNA, and Mt-rRNA), for each dFORCE biological replicate. **(g)** Z-score distribution in each experiment calculated after normalising the poly(A) tail lengths to the mean value of the reads from control genes. **(h)** Percentage of non-polyadenylated co-transcriptional reads (pre-mRNA$^{e}$) (blue) and endogenously polyadenylated reads (pre-mRNA$^{a}$) (red) identified in each dFORCE replicate.

### Figure S2

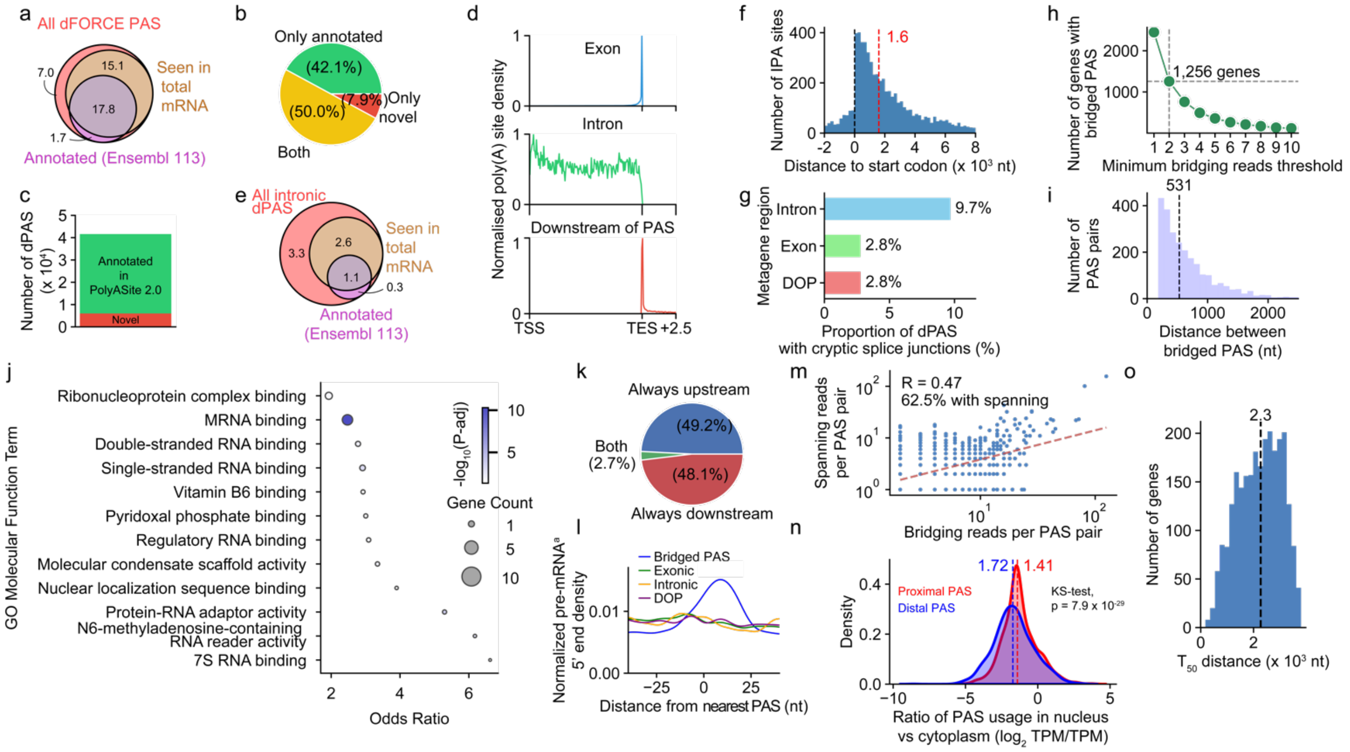


**Fig. S2. Supplemental information for dFORCE pre-mRNA 3' profiling. (a)** Venn diagram illustrating overlap between dFORCE poly(A) sites (PAS), total mRNA-supported PAS, and annotated PAS (see methods), for 4.2 x $10^4$ PAS identified in HeLa pre-mRNA[a]. Numbers represent PAS counts (in thousands). **(b)** Pie chart showing proportions of genes containing only annotated PAS (green), only novel PAS (red), or both types (yellow), for 1.2 x $10^4$ protein-coding genes. Percentages indicate fraction of total genes analysed. Classification of 'annotated' PAS corresponds to those in panel (a). **(c)** Stacked bar plot indicating the proportion of dFORCE clusters in (a) overlapping (green) or not overlapping (red) annotated poly(A) clusters from PolyASite 2.0 atlas (extended ±20 nt). **(d)** Metagene distribution of PAS location across genes, faceted by metagene (exons, introns, and downstream-of-PAS) regions. Coordinates are scaled from transcription start site (TSS; 0) to transcription end site (TES; 1.0), with PAS located downstream of TES mapped to an extended region (1.0–1.3). Each panel represents exon (top), intron (middle), and downstream-of-gene (DOG; bottom) categories. **(e)** Venn diagram illustrating overlap of intronic PAS, similar to (a), for 7.3 x $10^3$ intronic PAS. **(f)** Location of intronic PAS in (e) relative to start codons of expressed transcripts. Dashed red line indicates median distance (1.6 x $10^3$ nt). **(g)** Prevalence of PAS that present pre-mRNA alignments displaying cryptic splice junctions, stratified by PAS metagene regions (intronic, exonic, or DOP). Cryptic splice junctions are defined as those using any splice site at least 5 nt distant from all splice sites annotated in GRch38 (Ensembl release 113). **(h)** Count of protein-coding genes (y-axis) harbouring at least one pair of PAS bridged by varying minimum numbers of pre-mRNA[a] alignments (x-axis). Minimum threshold (≥2 bridging alignments), corresponding to 1,256 genes, is indicated on the plot. **(i)** Signed genomic distance between 2,495 pairs of bridged poly(A) sites in (h). Median genomic distance (531 nt) is indicated by the dashed vertical line. **(j)** Gene Ontology (GO) enrichment of molecular functions associated with genes presenting bridged poly(A) sites. Top 12 significant ($p$-adj < 0.05) Molecular Function (MF) GO terms (levels 3–5; ranked by odds-ratio) are shown for 1,256 genes with bridged PAS, compared to a background of 11,637 genes on which PAS were identified. **(k)** Venn diagram indicating role of bridged poly(A) sites, for 3,607 bridged poly(A) sites identified in (h-j). 'Always upstream' sites are only observed as being bridged to a downstream site, 'always downstream' sites are only observed as being bridged to an upstream site, and 'both' are observed as being bridged to both upstream and downstream poly(A) sites. **(l)** Metaplot showing the normalised distribution of pre-mRNA[a] alignment start positions relative to closest PAS, stratified by PAS category (bridged upstream PAS in any region, n = 1,872; exonic, n = 8,360; intronic, n = 1,178; or DOP, n = 2,671). Distributions are smoothed by a Gaussian filter ($\sigma$ = 4 nt). **(m)** Scatter plot depicting the relationship between number of bridging reads (x-axis) and spanning reads (y-axis) per polyadenylation site pairs, for 2,495 PAS pairs bridged by at least 2 alignments. Spanning reads are defined as those that end at the distal PAS but are uncleaved (i.e. span) the proximal PAS. 62.5% of PAS pairs shown are spanned by at least 1 read. **(n)** Kernel density estimates (KDE) illustrating the distribution of the $\log_2$-transformed ratio of polyadenylation site usage between dFORCE and total RNA for bridged PAS classified as proximal (red; n = 1,774) vs distal (blue, n = 1,724). Median values are indicated with dashed lines, and statistical significance between groups was assessed using the Kolmogorov–Smirnov test. **(o)** Histogram showing the $T_{50}$ distance (x-axis) for 2,570 protein coding genes where termination could be estimated.

### Figure S3

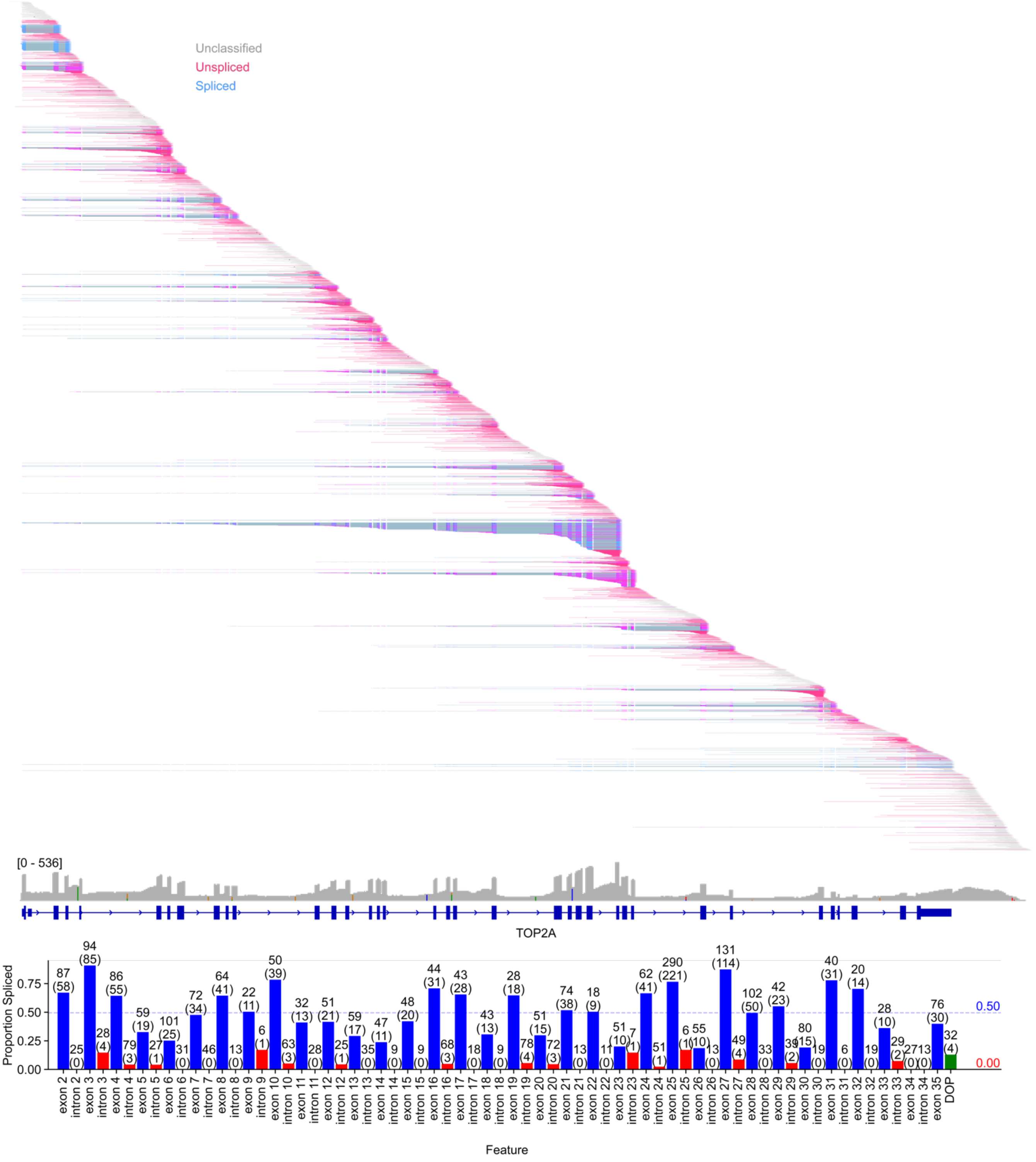


**Fig. S3. Splicing catalysis detected on *TOP2A* nonpolyadenylated pre-mRNA$^{e}$ reads** Top: dFORCE pre-mRNA$^{e}$ alignments at the *TOP2A* locus. Reads with at least 200 aligned bases that lacked cryptic splice junctions relative to the reference transcript are shown. Alignments are coloured by splicing classification and sorted by mapped 3' end coordinate. 'Spliced' reads contain ligated exon-exon junctions, unspliced reads contain at least one fully transcribed intron and no splice junctions, and all other reads, e.g. those fully contained in a single exon or intron, are classified as ambiguous. Bottom: Proportion of pre-mRNA$^{e}$ reads that are spliced (y-axis) ending at each of the *TOP2A* gene regions indicated in (a) (x-axis). Numbers above bars indicate the number of tested and spliced (in parenthesis) reads ending within each region. Dashed lines and values on right indicate median splicing efficiency across exons (blue) and introns (red).

### Figure S4

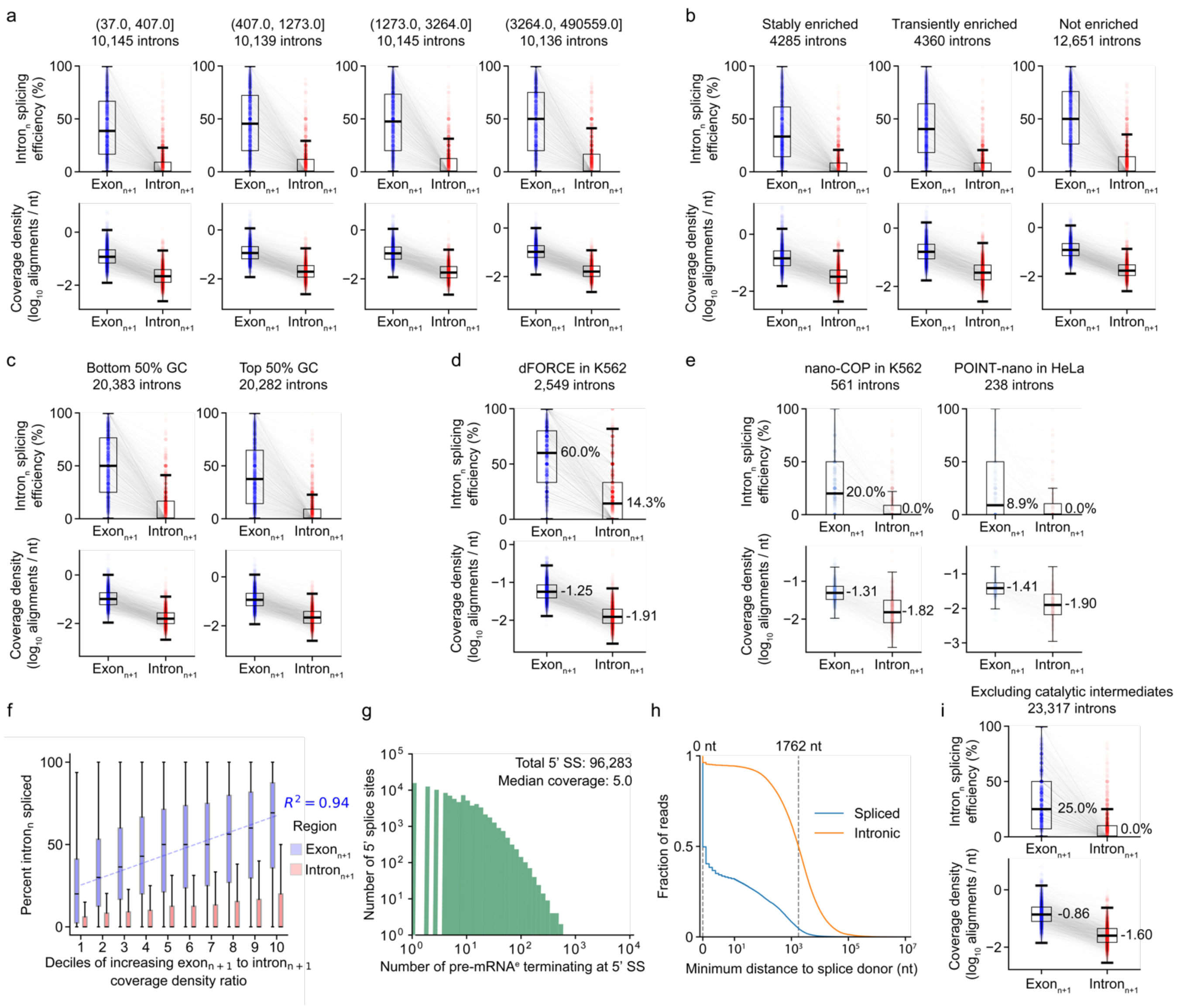


**Fig. S4. Supplemental information for exon$_{n+1}$ versus intron$_{n+1}$ splicing comparison. (a-c)** Splicing comparisons across subsets of introns in Fig. 3b, based on **(a)** quartiles of intron$_n$ length, **(b)** Association of genes with nuclear speckles in HeLa cells based on classifications from [1], and **(c)** intron$_n$ GC content. **(d)** Intron$_n$ splicing comparison in the style of (a-c) based upon a single biological replicate of dFORCE in K562 cells. **(e)** Splicing comparisons in the style of (a-d) based on nano-COP in K562 cells[2] and POINT-nano in HeLa cells[3] (see Table S1). **(f)** Stratification of 40,565 trios in Fig. 3b, based on deciles of increasing $\log_2$ ratio of alignment density in the exon$_{n+1}$ divided by alignment density in intron$_{n+1}$. Trendline measures the median extent of spliced reads ending in exon$_{n+1}$. **(g)** Distribution of number of unique 5' splice sites (y-axis) based on supporting number of CI* alignments ending within 1 nt of those splice sites (x-axis, $\log_{10}$ scale) **(h)** Distances between co-transcriptional pre-mRNA$^e$ alignment 3' ends and active 5' splice sites (5' SS), separated across alignments that contain any splice junctions ('spliced', n = 7.0 x $10^5$ alignments) and alignments that do not contain any splice junctions but span introns ('intronic', n = 8.0 x $10^6$ alignments), based on HeLa dFORCE. Median distances for spliced and intronic alignments are indicated on the plot. Alignments that are alternatively spliced compared to our annotation are omitted from this analysis. **(i)** Corresponding to Fig. 3d; paired intron-exon-intron splicing test, omitting CI* alignments ending within 1 nt of a splice donor.

### Figure S5

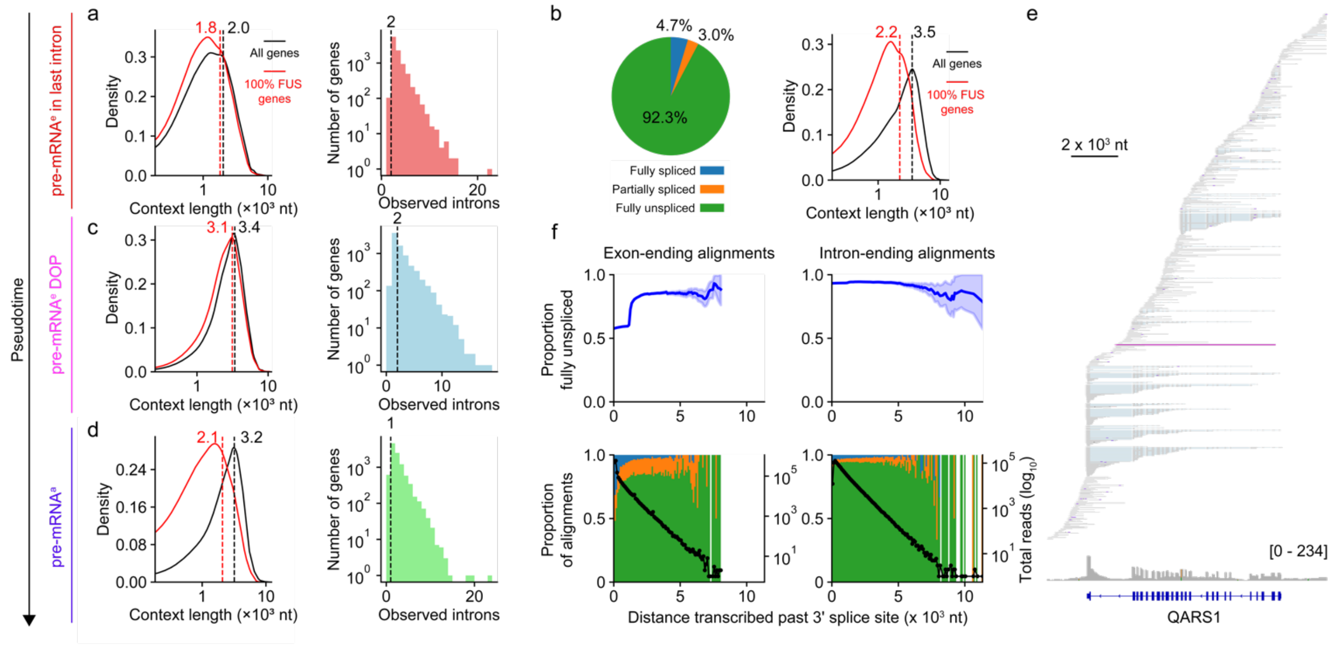


**Fig. S5. Supplemental information for splicing timing versus transcription elongation and 3' end formation. (a)** (Left) Distribution of context length per gene for 9,825 protein-coding genes where splicing of pre-mRNA$^{e}$ ending in the last intron was assessed (black curve) and the subset of these genes (n = 7,117) where 100% of pre-mRNA$^{e}$ in the last intron was fully unspliced (red curve). A total of 2.7 x $10^4$ introns were observed. Context length corresponds to the maximum aligned distance spanned by pre-mRNA$^{e}$ ending in each window for each gene. Dashed vertical lines represent the median context length for each set. (Right) For all genes tested in the left panel, the distribution of the highest number of observed introns spanned for per gene is shown. Dashed vertical line represents the median number of introns observed per gene. **(b)** (Left) For 10,599 genes, we show splicing classifications for all assessed pre-mRNA$^{e}$ which end in any intron except the first or last intron. (Colour scale: green, fully unspliced; orange, partially spliced; blue, fully spliced). Note, this classification only considers upstream introns and ignores the splicing of the intron in which the alignment ends, which is partially transcribed. (Right) Similar to (a), we show the context length distribution across all genes where internal intron-ending pre-mRNA$^{e}$ were tested (black, n = 10,599 genes) and for the subset of these genes where every internal intron ending read was fully unspliced (red, n = 3,258 genes). **(c)** Similar to (a), for pre-mRNA$^{e}$ ending downstream of poly(A) sites. 7,526 genes are plotted (black), of which 4,536 present exclusively fully unspliced alignments (red). A total of 1.7 x $10^4$ introns were observed. **(d)** Similar to (a), (c), for polyadenylated pre-mRNA$^{a}$ ending in last exons or downstream of the primary poly(A) site. A total of 2.0 x $10^4$ introns were observed. **(e)** IGV browser views of single-molecule dFORCE reads at *QARS1* locus (approximate GRCh38 coordinates chr3:49,093,900-49,107,000). All minus strand primary dFORCE alignments are displayed, sorted by 3' end position across the transcription unit. Alignment coloured in purple (read ID: 6e1da6dd-6fae-45a8-8da3-24b8ea85506f, 7 kb in length) is a pre-mRNA$^{e}$ molecule (poly(A) tail length of 10 nt) that spans all 23 introns in the gene. **(f)** Extent of splicing versus distance transcribed past 3' splice site, binned by exon-ending pre-mRNA$^{e}$ (left panels) and intron-ending pre-mRNA$^{e}$ alignments (right panels). (Top panels) Smoothed proportion of fully unspliced reads versus extent of transcription past the 3' splice site, binned in 100 nt intervals. Shaded band indicates the 95% CI derived from the binomial variance within each rolling window, smoothed across 20 bins. (Bottom panels) Stacked bar plot (left y-axis) displaying the proportion of fully unspliced, partially spliced and fully spliced alignments within each bin indicated in the top panel. Colours correspond to panel (b). Overlaid black line (right y-axis) indicates the number of alignments tested within each bin. X-axis (shared) corresponds to the distance between the 3' splice site and the alignment 3' end. Note, CI* are not excluded from exon or intron-ending pre-mRNA$^{e}$.

### Figure S6

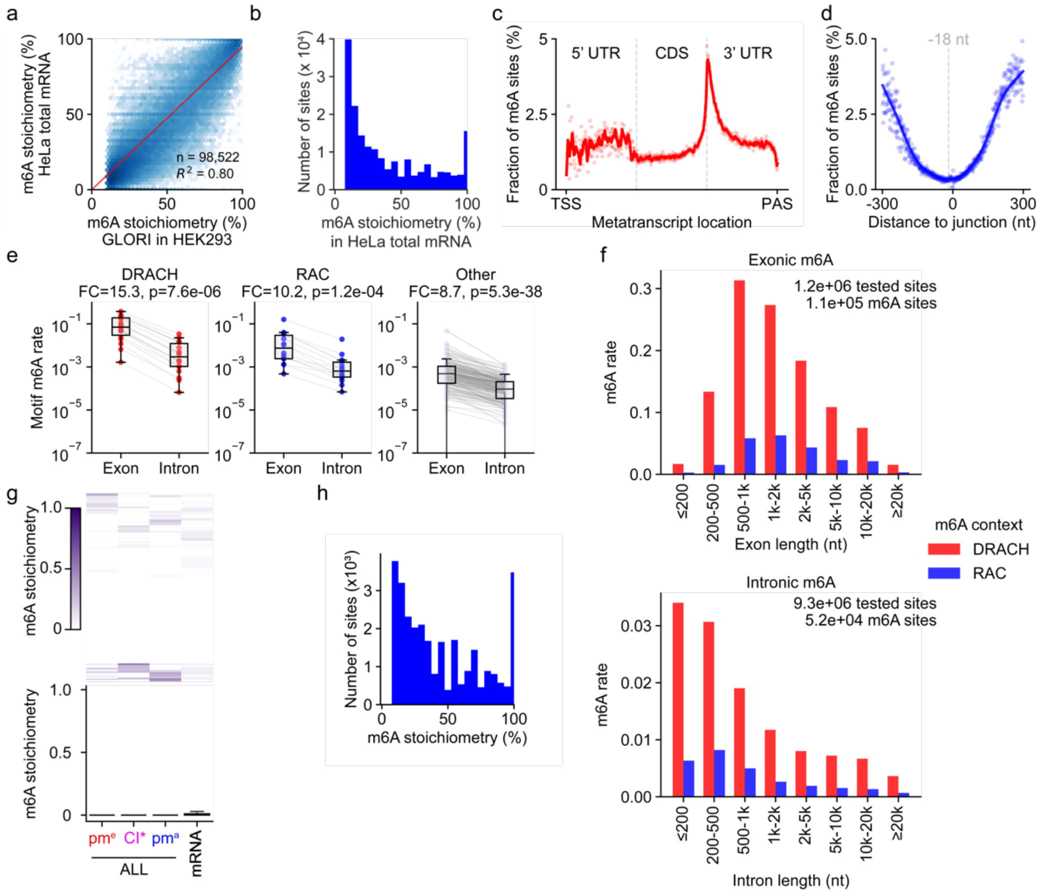

**Fig. S6. Supplemental information for m6A detection and timing in HeLa cells. (a)** Validation of nanopore m6A site discovery by comparison between m6A sites detected by nanopore in HeLa total mRNA (x-axis) compared to published m6A levels identified on HEK293 total RNA by GLORI[4]. **(b)** Distribution of site-level m6A stoichiometry for 182,261 m6A sites detected from total mRNA aligned to the GRCh38 genome. **(c)** Metatranscript profile of m6A rates (i.e. proportion of modified A sites over number of A nucleotides in each metatranscript bin) based on total mRNA aligned to Ensembl 113 (GRCh38) transcript isoforms. Transcripts are divided into 600 bins, equally divided across 5'UTR, CDS, and 3'UTR regions (scatterplot). A locally weighted scatterplot smoothing (lowess) trendline (span = 2 metajunction bins) is fitted to the data. **(d)** Metajunction profiles of m6A rate (proportion of modified A sites over tested A sites in each metajunction bin) based on distances between tested sites and the closest annotated splice donor in Ensembl 113 (GRCh38) (Scatterplot). A lowess trendline (span = 2 nucleotides) is fitted to the data. **(e)** m6A methylation rate (y-axis, $\log_{10}$) in introns versus exons (x-axis), stratified by DRACH (left), non-DRACH RAC (centre) or non-RAC (right) sequence contexts. Motif m6A rate is defined as the number of tested instances of that motif in genomic space, divided by the number of methylated instances of that motif. **(f)** Pre-mRNA m6A methylation rate (y-axis) on exons (upper panel) and introns (lower panel), binned by exon or intron length (x-axis) and stratified by sequence context: DRACH (red), non-DRACH RAC (blue). **(g)** m6A dynamics of silenced strong DRACH sites. Testing n = 21,164 instances of the most frequent m6A-positive 5-mers (GGACT, GGACA or GAACT), that were silenced in total mRNA (<5% m6A level) and sequenced by at least 5 reads in all fractions shown. Across pre-mRNA[a], CI*, and pre-mRNA[e] fractions, 93-97% of the tested sites displayed <5% m6A stoichiometry, 99.9% <30% (Fig. 5s). Thus, sites destined to remain unmethylated in mature transcripts are typically already repressed in pre-mRNA across all maturation steps. **(h)** Distribution of site-level m6A stoichiometry for 2.9 x $10^4$ m6A sites detected on CI* alignments aligned to GRCh38 genome.

### Figure S7

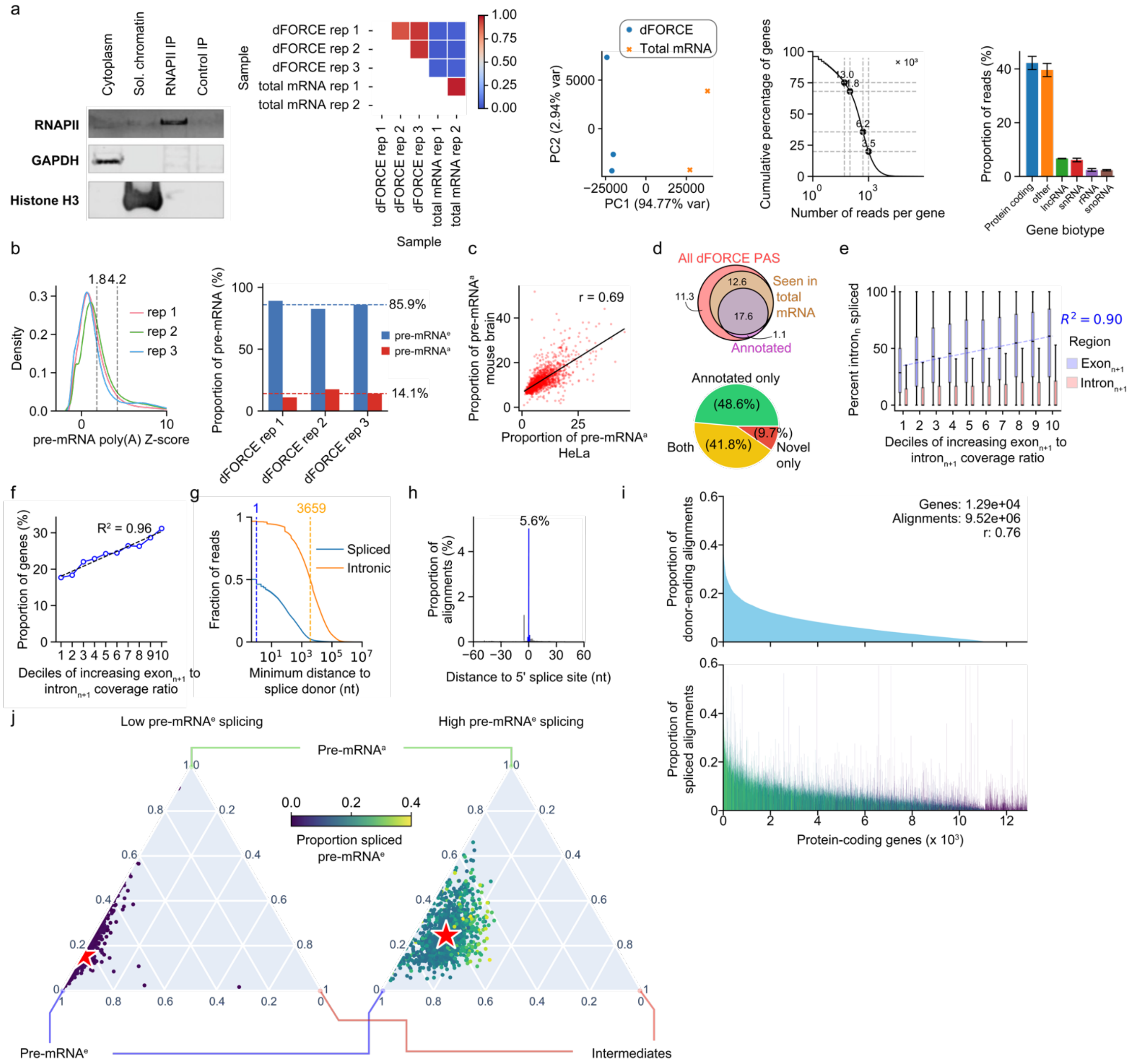

**Fig. S7. Supplemental information for *in vivo* dFORCE. (a)** (Left) Western blotting against RNAPII CTD, GAPDH and Histone H3 antibodies for mouse brain, cytoplasm, soluble chromatin, RNAPII CTD IP and control antibody IP (anti Drosophila Aubergine). 0.3%, 2.5%, and 25% of the cytoplasmic, the soluble chromatin, and the IP fractions were loaded. (inside-left and centre panels) Correlation of gene expression levels between mouse brain dFORCE and total mRNA sequencing, and PCA representing the same information, for 12,033 protein-coding genes. (centre-right) Distribution of primary alignments per protein-coding gene for 1.7 x $10^7$ dFORCE primary alignments mapping to 1.7 x $10^4$ protein-coding genes. Number of genes with at least 50, 100, 500, or 1000 reads are labelled on the plot. (right) Distribution of gene biotypes from primary alignments of mouse dFORCE reads against GRCm39 (Ensembl release 113, unfiltered annotation). Error bars indicate standard error of the mean across 3 biological replicates. **(b)** (Left) Distributions of normalised pre-mRNA poly(A) z-scores for 3 dFORCE replicates. Maximum z-score limit for pre-mRNA$^e$ (1.8) and minimum limit for pre-mRNA$^a$ (4.2) are labelled on the plot. (Right) Proportion of co-transcriptional (pre-mRNA$^e$; blue) and post-transcriptional (pre-mRNA$^a$; red) per library for poly(A)-classified reads across 3 dFORCE replicates. **(c)** Proportion of pre-mRNA$^a$ per gene between HeLa (x-axis) and mouse brain (y-axis) for 1,495 orthologous gene pairs. **(d)** (Top) Classification of mouse brain dFORCE poly(A) site clusters for 48,176 identified clusters. 'Annotated' clusters are those within 20 nt of transcript end site annotated in Ensembl 113. Values shown are x$10^3$. (Bottom) Novelty classification of PAS for 13,014 genes. 'Annotated' PAS correspond to those in the left panel. **(e)** Stratification of 32,027 trios in Fig. 6c based on deciles of increasing $\log_2$ ratio of alignment density in the exon$_{n+1}$ divided by alignment density in intron$_{n+1}$. Trendline measures the median extent of spliced reads ending in exon$_{n+1}$. **(f)** Based on the same stratification as (k), the proportion of genes in each bin with any efficiently spliced introns (intron$_n$ at least 50% spliced). **(g)** Distances between pre-mRNA$^E$ alignment 3' ends and active 5' splice sites (5' SS), separated across alignments that contain any splice junctions ('spliced', n = 3.8 x $10^5$ alignments) and alignments that do not contain splice junctions but span introns ('Intronic'; n = 9.2.x $10^6$ alignments). Median values for spliced, and non-spliced reads are indicated on the plot. Alignments that are alternatively spliced compared to our annotation are omitted. **(h)** Number of pre-mRNA$^e$ reads with 3' ends at various distances to active splice donors. 5.6% of mouse brain pre-mRNA$^e$ end within 1 nt of an active splice donor. **(i)** Top: Ranking (x-axis) of protein-coding genes by the proportion of pre-mRNA$^e$ reads end within 1 nt of a 5' splice site, classified as catalytic intermediates (y-axis). Bottom: In the same ranking, proportion of pre-mRNA$^e$ alignments that display a splice junction (y-axis) across the same set of genes. Genes are coloured by the proportion of spliced reads that end within 1 nt of a 5' splice site, compared to the number of spliced reads ('Donor ending spliced alignments). **(j)** Ternary plot representing the proportion of pre-mRNA[a], pre-mRNA[e] (excluding donor-ending reads) and catalytic intermediates (i.e. donor-ending pre-mRNA[e] reads) for protein-coding genes with at least 50 alignments and up to 40% spliced pre-mRNA[e]. Left panel: bottom 10% of genes according to spliced pre-mRNAe. Right panel: top 10% of genes. Red stars indicate the average proportions of (non-intermediate) pre-mRNA[e], catalytic intermediates, and post-transcriptional pre-mRNA[e] in each gene set. 1 x $10^3$ genes are displayed per panel. Middle colour scale indicates the proportion of pre-mRNA[e] reads that are spliced in each panel.

### Figure S8

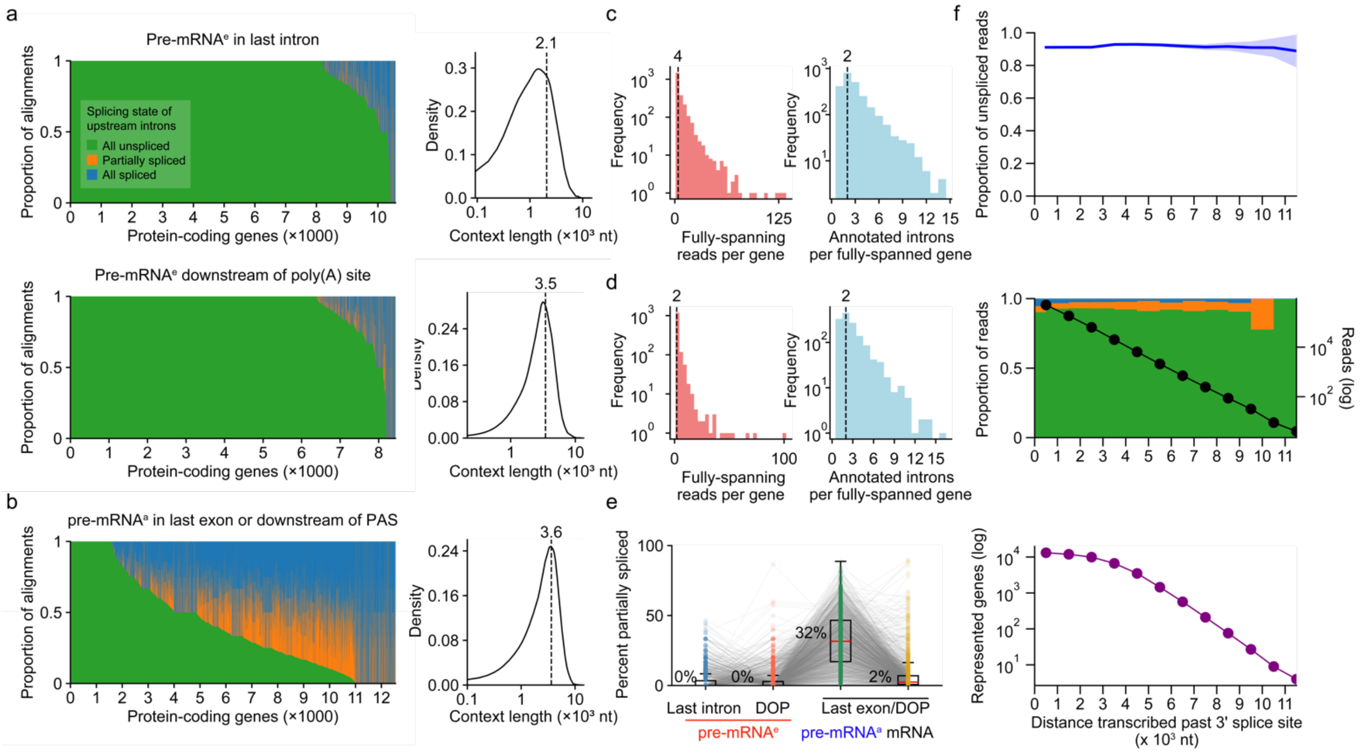


**Fig. S8. *In vivo* timing of splicing catalysis. (a-b)** Left panels: Distribution of spliced, unspliced and partially-spliced alignments over (top panel) pre-mRNA$^e$ ending in the last intron of 10,575 protein-coding genes, (middle-panel) pre-mRNA$^e$ ending downstream of the primary poly(A) site of 8,438 protein-coding genes, and (bottom panel) pre-mRNA$^a$ ending at the 3' end (last exon or downstream of poly(A) site) of 12,456 genes. A total of 3.0 x $10^4$ (top), 2.0 x $10^4$ (middle) or 2.8 x $10^4$ distinct unspliced introns were observed in each transcriptional window. Colour scale per gene: Green: fully unspliced alignments, orange: partially spliced alignment, blue: fully spliced alignment. Note, the splicing classification only considers upstream introns and ignored the last intron, which is partially transcribed. Right panels: Density plots indicating the context length across each set of genes in the left panel, within the region specified. **(c-d)** Abundance of completely-unspliced reads (spanning all introns in the gene) per protein-coding gene (left) and number of introns on completely unspliced genes (right) for 2,388 genes completely spanned by pre-mRNA$^e$ (top) and 1,403 genes completely spanned by pre-mRNA$^a$ (bottom). **(e)** Proportion of partially-spliced alignments compared to partially-spliced, fully-spliced and fully unspliced, for 1,445 protein coding genes with at least 5 assessed alignments in all of pre-mRNA$^e$ in the last intron, pre-mRNA$^e$ downstream of the poly(A) site, pre-mRNA$^a$ in the last exon or downstream of the poly(A) site, and total mRNA ending in the last exon or downstream of the poly(A) site. **(f)** Proportion of fully unspliced RNA as a function of distance transcribed past the 3' splice site. Alignments ending within exons were excluded. Top panel: Smoothed proportion of fully unspliced reads per length bin, with shaded regions representing the 95% confidence interval (CI). Middle panel: Proportions of fully unspliced, partially spliced, and fully spliced reads per length bin (left y-axis), overlaid with the total number of reads tested per bin shown on a $\log_{10}$ scale (right y-axis). Bottom panel: Number of unique genes represented per length bin ($\log_{10}$ scale).

### Figure S9

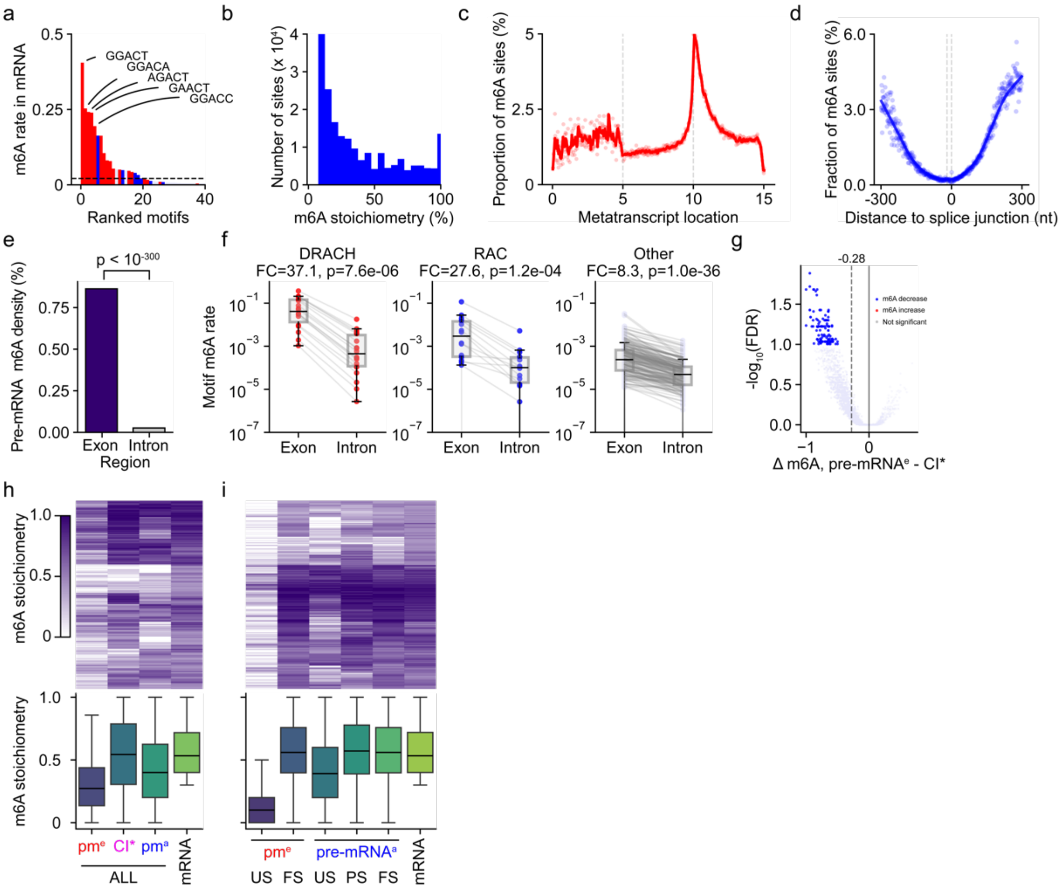


**Fig. S9. *In vivo* dynamics of m6A methylation. (a)** Methylation rate across top 40 A-centred 5mer sequences from total mRNA mouse brain total mRNA genomic alignments. Motifs are ranked by m6A methylation rate (y-axis), defined as the number of genomic instances of that motif classified as m6A sites (≥5% m6A stoichiometry and ≥2 m6A-modified reads) as a proportion of the total number of genomic instances that were tested in total mRNA. Bars are coloured by motif type. Red corresponds to DR<u>A</u>CH motifs, blue to non-DRACH R<u>A</u>C motifs, and grey to other motif types. **(b)** Distribution of site-level m6A stoichiometry for 2 x $10^6$ m6A sites detected from mouse brain total mRNA aligned to GRCm39 genome. **(c)** Metatranscript profile of m6A rates (i.e. proportion of modified A sites over number of A nucleotides in each metatranscript bin) based on total mRNA aligned to GRCm39 transcript isoforms. Transcripts are divided into 600 length bins, equally divided across 5'UTR, CDS, and 3'UTR regions (scatterplot). A lowess trendline (span = 2 metajunction bins) is fitted to the data. **(d)** Metajunction profiles of m6A rate (proportion of modified A sites over tested A sites in each metajunction bin) based on distances between tested sites and the closest annotated splice donor in GRCh38 (Scatterplot). A lowess trendline (span = 2 nucleotides) is fitted to the data. **(e)** Density (y-axis) of m6A sites across pre-mRNA exons and introns. Density was defined as the number of m6A positive genomic sites divided by the number of tested genomic A sites. Chi-square test was used. **(f)** m6A methylation rate (y-axis, $\log_{10}$) in introns versus exons (x-axis), stratified by DRACH (left), non-DRACH RAC (centre) or non-RAC (right) sequence contexts. Motif m6A rate is defined as the number of tested instances of that motif in genomic space, divided by the number of methylated instances of that motif. **(g)** Differential m6A methylation test for 4.9 x $10^3$ total mRNA m6A sites detected to be at least 30% methylated in dFORCE pre-mRNAe or CI* and spanned by at least 5 reads in both fractions. x-axis: Difference in site stoichiometry between pre-mRNA$^e$ – catalytic intermediates, y-axis: FDR-adjusted p-value. Vertical dashed line indicates the median stoichiometry difference across the sites; Sites hypomethylated on pre-mRNAe (FDR ≤ 0.1 and Δm6A ≥ = -0.1) are coloured in blue, and those hypermethylated on pre-mRNAe (FDR ≤ 0.1 and Δm6A ≥ = 0.1) are coloured in red. **(h)** Top: Heatmap of m6A stoichiometry values for 1,702 m6A sites which are constitutive in total mRNA (≥ 30% stoichiometry) which are tested in at least 5 alignments in total RNA and each dFORCE fractions. Bottom: Boxplots representing the stoichiometry distribution in the same fractions. pm$^e$: pre-mRNAe, CI*: splicing catalytic intermediates, pm$^a$: pre-mRNA$^a$. **(i)** Similar to (j), testing n = 1,735 m6A sites that were detected as constitutive in total mRNA and tested in at least 5 alignments in all fractions shown. Pre-mRNA fractions are stratified by splicing status (US: unspliced, PS: partially spliced, FS: fully spliced).

### Figure S10

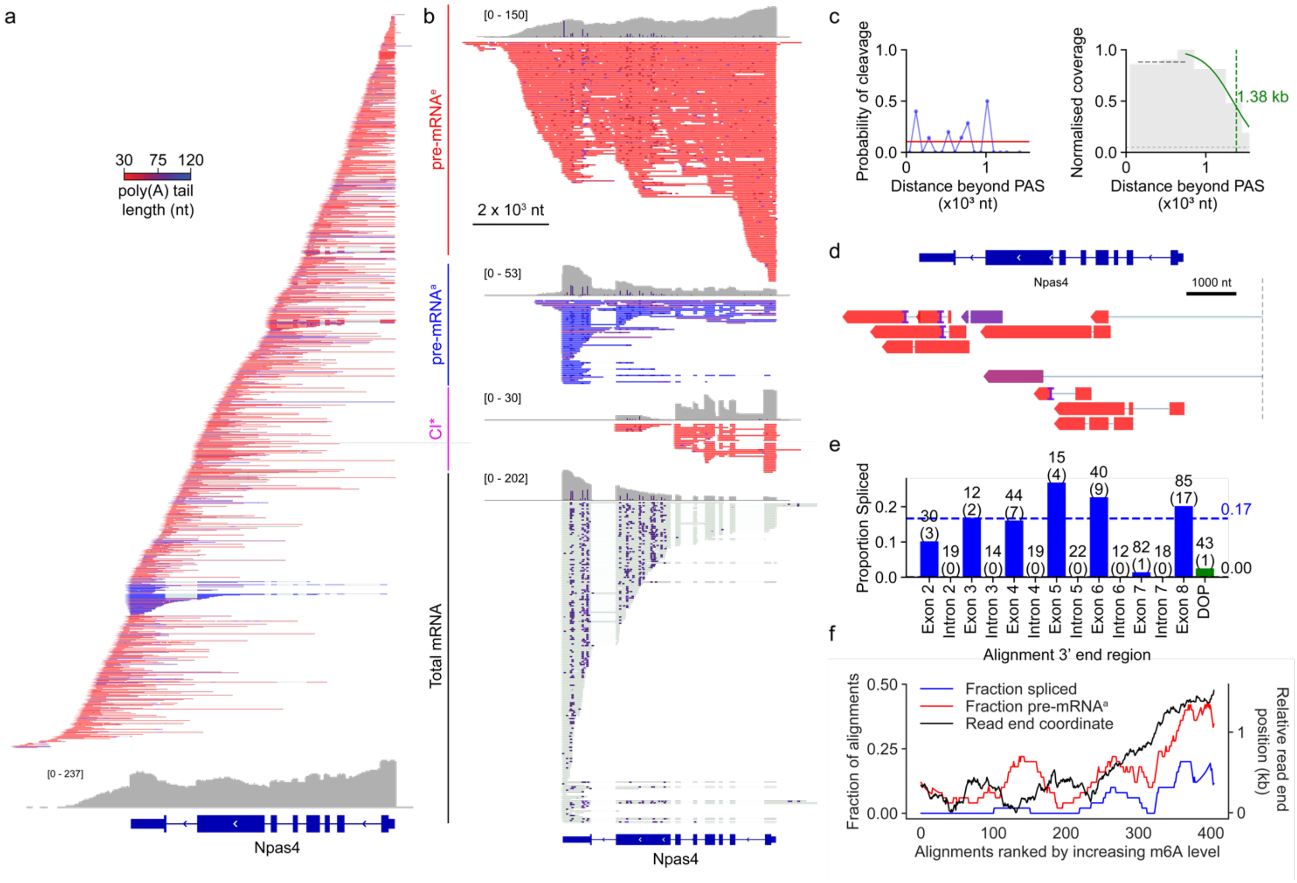


**Fig. S10. Integrated processing of Npas4 *in vivo*. (a)** IGV browser views of single-molecule dFORCE reads at Npas4 locus from dFORCE in the mouse brain (GRCm39 approximate region: chr19:5,032,000-5,042,000). Alignments are coloured by poly(A) tail length and sorted by alignment 3' ends, representing RNA polymerase II position, across the transcription unit. All minus strand primary alignments are shown without omission. Total coverage over gene regions and annotated gene structures are indicated at the bottom of the figure. Colour scale corresponds to single-molecule poly(A) tail length. **(b)** dFORCE (top 3 panels) and total mRNA (bottom panel) m6A-basecalled alignments at the mouse *Npas4* locus. dFORCE alignments are coloured by poly(A) tail length and separated into pre-mRNA$^{e}$, pre-mRNA$^{a}$ and splicing catalytic intermediate (CI*) fractions. Only positive strand alignments are displayed. Alignments are not sorted by 3' end position. Colour scale for dFORCE poly(A) tail lengths corresponds panel (a). **(c)** (Left) Coverage fraction downstream of PAS shown in 200 nt bins, normalized to peak coverage. Logistic regression (green line) is fitted to the coverage decline; dashed green line marks the position where coverage falls to half the median peak fraction. (Right) Binned cleavage probabilities at Npas4 (blue circles) fitted with a logistic regression (red line). **(d)** Cryptically spliced alignments at Npas4 locus (n = 10). These alignments are removed before classifying splicing. Colours correspond to panel (a). **(e)** Proportion of pre-mRNA$^{e}$ reads that are spliced (y-axis) ending at each of the Npas4 gene regions indicated on the x-axis. Numbers above bars indicate the number of tested and spliced (in parenthesis) reads ending within each region. Dashed lines and values on right indicate median splicing efficiency across exons (blue) and introns (red). **(f)** Splicing, polyadenylation and read-end position properties of *Npas4* pre-mRNA alignments ranked by proportion of m6A sites (identified in total mRNA) that are m6A positive on the dFORCE alignments.

### Supplemental materials

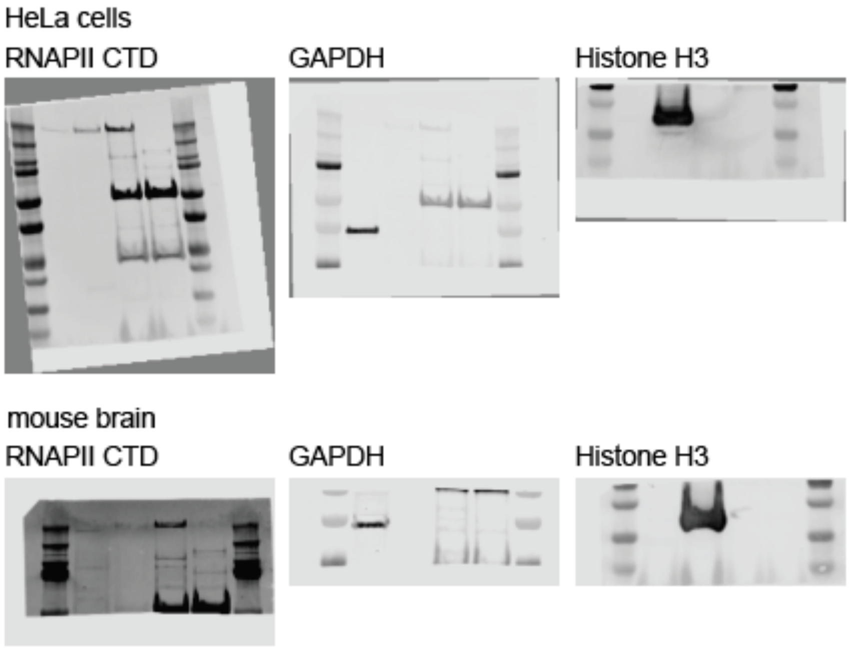


Shown are raw images of the western blotting presented in Fig. S1a and S7a.
